# Transcriptional activity generates chromatin motion that drives nuclear blebbing

**DOI:** 10.1101/2025.05.20.655131

**Authors:** Kelsey Prince, Katie Lin, Andy Li, Nick Borowski, Andrew D. Stephens

## Abstract

Abnormal nuclear morphology is a hallmark of human diseases, including cancers and age-related disorders. Previously, maintenance of nuclear morphology and integrity was thought to be solely dependent on a force balance between nuclear mechanical resistance and actin antagonism. However, our recent work revealed that inhibiting RNA polymerase II suppresses nuclear blebbing independent of altering force balance, but the mechanism remains unknown. Through removing cell culture media serum and then adding it back, we can decrease and then restore transcriptional activity. Decreasing transcriptional activity decreases nuclear bleb formation, stability, and rupture while returning transcriptional activity restores nuclear blebbing. These modulations of transcriptional activity did not alter nuclear or actin mechanics. The mean square displacement (MSD) of chromatin domains labeled via transfected Cy3-dNTPs revealed that transcription activity regulates chromatin motion. To determine if increasing chromatin motion is a mechanism to increase nuclear blebbing, we used an established RAD51 inhibitor BO2. We verified BO2 increases chromatin domain motion which resulted in increased nuclear blebbing. We reveal the mechanism by which transcriptional activity drives nuclear blebbing is through chromatin motion. Thus, two hallmarks of human disease are directly linked via transcriptional activity and abnormal nuclear shape.

**Statement of Significance:**

- Nuclear blebs are hallmarks of disease progression that cause dysfunction, but how they are formed remains unanswered.
- We find that chromatin motion generated by transcriptional activity is essential for both nuclear bleb formation and stability. This was independent of changes in nuclear stiffness or actin antagonism.
- This finding provides a key advancement in our understanding of nuclear bleb formation. Furthermore, it reveals transcriptional activity as a novel contributor to nuclear blebbing in addition to the paradigm of nuclear shape determined as a force balance between nuclear resistance and actin antagonism.

## Introduction

Nuclear blebbing is a phenomenon associated with human diseases, including cancer and age-related disorders such as progeria. Nuclear blebbing is a type of abnormal nuclear deformation that presents as a herniation of the nucleus > 1 µm in size and has decreased DNA density (Bunner *et al*., 2024; Pujadas Liwag *et al*., 2025). Understanding the mechanisms underlying nuclear bleb formation may aid in the treatment or prevention of human diseases. Previous research has shown that nuclear shape maintenance is determined by a force balance between chromatin and lamins that provide nuclear resistance to actin antagonism via confinement and contraction (Hatch and Hetzer, 2016; Stephens *et al*., 2018; Mistriotis *et al*., 2019; Kalinin *et al*., 2021; Vahabikashi *et al*., 2022; Pho *et al*., 2023; Manning *et al*., 2025). Independent of nuclear mechanics, transcription inhibition has been shown to decrease nuclear blebbing (Berg *et al*., 2023), though the mechanism remains unknown.

We aim to elucidate the role of transcription activity in nuclear blebbing and rupture. Serum starvation, removal of most serum from the cell media, is well-documented approach for decreasing transcriptional activity (Kirkconnell *et al*., 2016). Serum stimulation, adding serum back the media, restores the transcriptional activity of serum starved cells in a matter of hours (Tullai *et al*., 2007; Kirkconnell *et al*., 2016, 2017). This method decreases then returns transcriptional activity in the same cells providing a high level of control.

The prevailing hypothesis is that transcriptional motor activity contributes to chromatin movement that pushes against the nuclear envelope to drive nuclear blebbing. An alternative hypothesis is that changes in transcription activity cause off-target effects on known nuclear and actin force balance components. Support for the chromatin motion hypothesis is that RNA polymerase II (RNA Pol II) is molecular motor that walks along the DNA during transcription exerting a pulling force in the direction of travel and pushing in the opposite direction. The motor activity of RNA Pol II can provide 10 piconewtons of force (Herbert *et al*., 2008). Physics modeling suggests only a fraction of RNA Pol II’s force capability is required to generate chromatin motion that can alter nuclear shape via collisions with the nuclear periphery (Liu *et al*., 2021; Berg *et al*., 2023). Previous studies have shown that transcriptional activity is important for micron-sized chromatin domain coherent motion (Zidovska *et al*., 2013; Shaban *et al*., 2018, 2020; Locatelli *et al*., 2022; Chu *et al*., 2024). Oppositely, transcriptional activity suppresses single nucleosome motion (Nagashima *et al*., 2019). BO2 is a RAD51 inhibitor shown to increase chromatin domain motion but not nucleosome movement (Maarouf *et al*., 2024). Thus, we can separate and modify chromatin domain motion independently from single nucleosome motion which behaves differently.

In order to test the hypothesis that transcription activity generates chromatin motion, cells can be transfected with Cy3-dNTPs to label chromatin replication domains (Albiez *et al*., 2006; Pabba *et al*., 2023). Chromatin motion can be quantified by mean squared displacement (MSD) (Berg, 2018). Furthermore, recently published work provides a manner to increase chromatin motion via use of RAD51 inhibitor BO2 (Maarouf *et al*., 2024). To test the alternative hypothesis, we can separately determine the nuclear rigidity provided by chromatin and lamins via dual micropipette micromanipulation force measurements (Stephens *et al*., 2017; Currey *et al*., 2022), which have been verified by other techniques (Hobson *et al*., 2020; Bergamaschi *et al*., 2024). The contribution of actin antagonism to nuclear shape via confinement can be measured via nuclear height and actin contraction measured by levels of phosphorylated myosin light chain 2 (pMLC2,(Pho *et al*., 2023)). Thus, there is an established suit of tools for determining a mechanism for transcriptional activity’s effect on nuclear shape.

To determine the role of transcriptional activity in nuclear blebbing, we used serum starvation and add back then measured the physical properties of the nucleus. Immunofluorescence of active RNA polymerase II and click-it chemistry for newly synthesized RNA confirmed serum starvation and add back respectively decreases and then restores transcriptional activity. Nuclear blebbing in population and time lapse imaging provide measurements of nuclear blebbing percentages and stability along with nuclear ruptures. We then measured actin and nuclear mechanics to determine if transcriptional activity modulations via serum were altering these established nuclear blebbing determinants. Mean squared displacement (MSD) of replication domains labeled by Cy3-dUTPs revealed changes in chromatin motion. Finally, we treat cells with RAD51 inhibitor BO2 to increase chromatin motion, independent of transcription activity, to determine the effect on nuclear blebbing. Overall, we find that transcription modulation via serum starvation and add back affects chromatin motion to drive nuclear blebbing.

## Results

### Transcriptional activity is decreased by serum starvation and restored by serum add back

We performed immunofluorescent imaging to assess the levels of active RNA Pol II markers via phosphorylation at Ser5 and Ser2, which correspond respectively to transcription initiation (Komarnitsky *et al*., 2000; Schwer and Shuman, 2011) and elongation (Barilla *et al*., 2001; Kim *et al*., 2004). For our control we used untreated (UNT) mouse embryonic fibroblasts (MEFs, **Figure 1A-C**). Valproic acid (VPA) is a histone deacetylase inhibitor established to increase euchromatin and transcriptional activity (Göttlicher *et al*., 2001). Treatment of cells with VPA revealed an increase in RNA Pol II pSer5 and pSer2 markers relative to untreated. An established inhibitor of transcription is alpha amanitin (AAM) which binds to and degrades RNA Polymerase II (Kedinger *et al*., 1970; Bensaude, 2011). Treatment of AAM revealed a decrease in RNA Pol II pSer5 and pSer2 markers relative to untreated. Thus, we can effectively measure changes in transcriptional activity via RNA Pol II pSer5 and pSer2.

**Figure 1.**
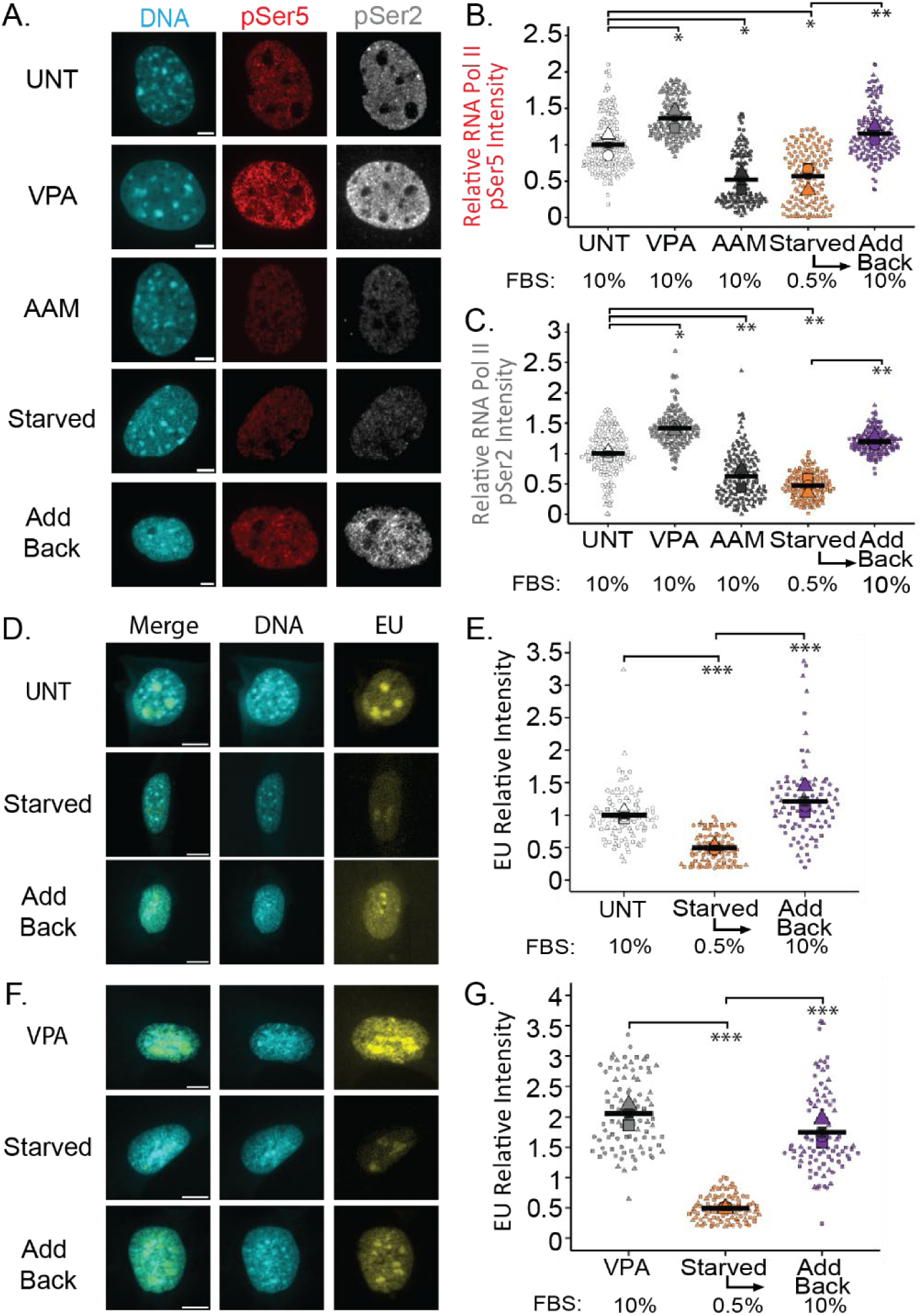
Transcriptional activity is decreased by serum starvation and restored by serum add back. (A) Example images and (B,C) graphs of relative fluorescence intensity of active RNA Pol II marking (B) initiation via pSer5 and (C) elongation via pSer2 in MEF cells untreated control (UNT), chromatin decompaction (VPA), RNA Pol II inhibition (AAM), serum starved, and serum add back. (D,F) Example images and (E, G) graphs of relative fluorescence intensity of EU click-it chemistry to measure new synthesized RNA in untreated, serum starved, and serum add back in (D,E) MEF WT and (F, G) MEF VPA-treated cells. For all conditions serum starvation was for 72 hours and serum add back was for 3 hours. N=3 for all data, >50 cells per replicate. Error bars represent standard error and statistical tests are one-way ANOVA with a post-hoc Tukey test, with significance denoted by *= p<0.05, **= p<0.01, and ***=p<0.001. Scale bar =10µm.

To modulate transcriptional activity, we altered levels of serum in the cell media. Based on previous work, serum starvation should lead to decreased transcriptional activity (Kirkconnell *et al*., 2016; Galves *et al*., 2023). Cells were starved for 72 hours by decreasing Fetal Bovine Serum (FBS) from the standard 10% to 0.5% in the cell culture media. Serum starvation significantly decreases transcriptional activity compared to untreated (UNT) mouse embryonic fibroblasts (MEFs, **Figure 1A-C**). The levels of transcriptional activity in serum-starved cells were similar to those observed upon transcription inhibition in AAM-treated cells. Thus, serum starvation provides a means to decrease transcriptional activity.

Transcriptional activity in serum starved cells can be restored in a few hours through adding back serum to the cellular media, we denote as add back (Kirkconnell *et al*., 2017). Specifically, after 72 hours of serum starvation in 0.5% FBS media, serum was added back to the media at the standard level of 10% FBS. Upon serum add back for 3 hours, transcriptional activity measured by active RNA Pol II pSer5 and pSer2 was restored to similar levels compared to untreated cells (**Figure 1A-C**), which was significantly higher than in serum starved cells. Serum add back, also called stimulation, for more than twelve hours significantly increased transcriptional activity at both initiation and elongation markers compared to untreated (**Supplemental Figure 1A-C**). To further corroborate these findings, we used a second cell line,

HT1080, a human fibrosarcoma model. Serum-starved HT1080 cells also exhibited decreased transcriptional activity which returned to wild type levels within 3 hours of serum add back and increased significantly after twelve hours (**Supplemental Figure 1D-F**). Taken together, serum starvation and add back provide a method to decrease then quickly restore transcriptional activity.

To recapitulate these findings of changes in transcriptional activity via a different readout, we measured newly synthesized RNA levels by adding the uridine analog 5-ethynyluridine (EU) for 1 hour and then performing Click-iT chemistry (Jao and Salic, 2008). Serum starvation led to a significant decrease in newly synthesized RNA levels (**Figure 1D-G**). Subsequently, serum add back restored newly synthesized RNA levels to untreated levels within three hours. This data confirms that transcriptional activity is effectively regulated by altering serum levels in the cell culture media.

### Transcriptional activity modulation via serum affects nuclear blebbing, rupture, and reabsorption

Once transcriptional activity changes were confirmed, the next step was to determine whether serum manipulation affected nuclear blebbing and rupture. Live-cell and time lapse imaging were used to quantify the percentage of nuclear blebs upon serum starvation and add back. Nuclear blebs were determined as a greater than 1 µm protrusion of the nucleus with decreased DNA density when compared to the nuclear body (**Figure 2A**).

**Figure 2.**
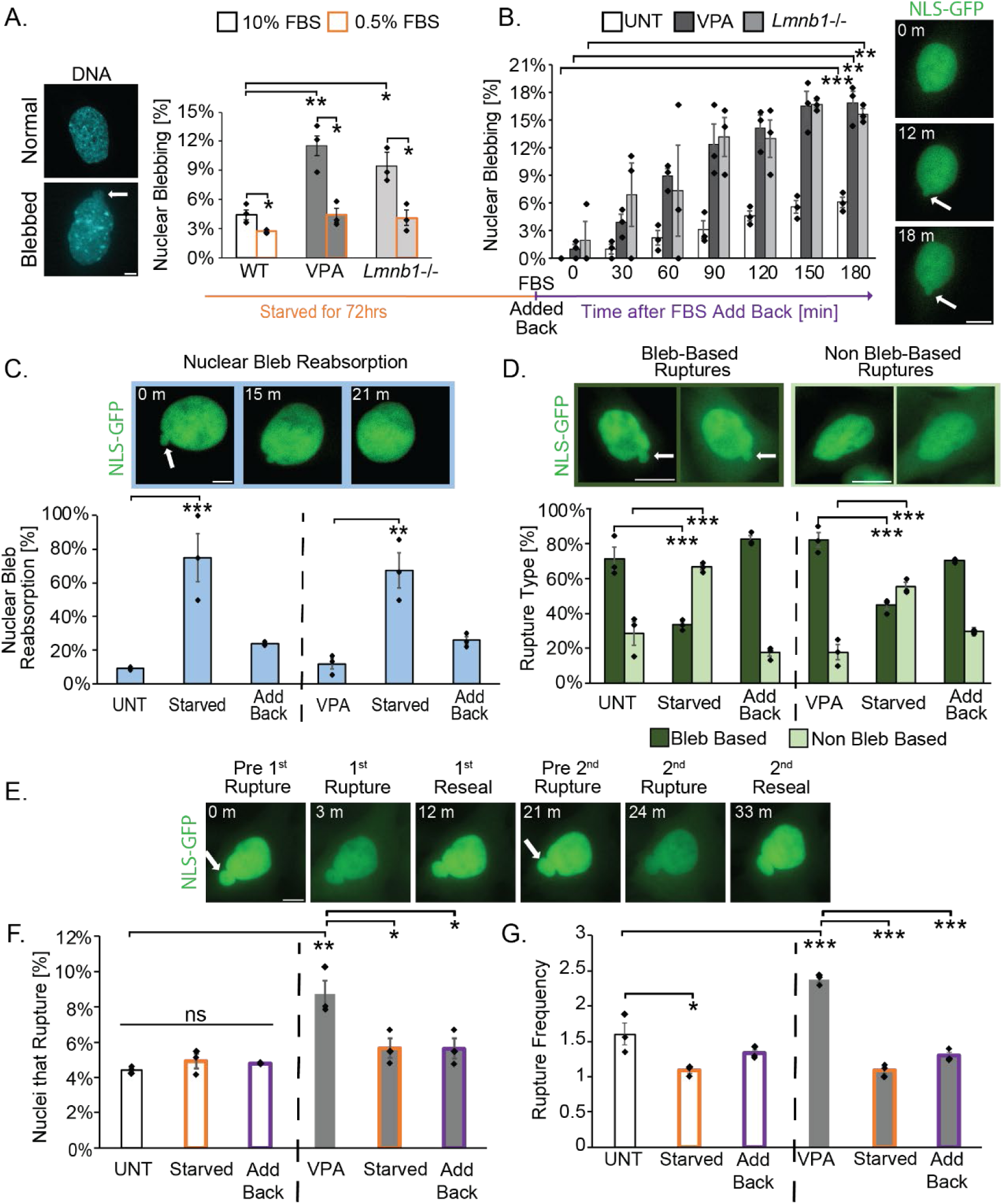
Serum manipulation regulates nuclear blebbing and rupture. (A) Example images of normal and blebbed MEF wild type nuclei stained with Hoechst and graph of blebbing percentage in normal (10% FBS, black outline) and serum starved (0.5% FBS, orange outline) in MEF cell conditions untreated (UNT), chromatin decompaction (VPA), and MEF *Lmnb1-/-*.(B) Graph and example images of bleb formation over 3-hour time lapse upon serum add back in cells previously starved for 72 hours in MEF wild type untreated (UNT), chromatin decompaction (VPA), and *Lmnb1-/-*. (C) Example images of a MEF nuclear bleb that ruptured and was reabsorbed (3-minute intervals) and graph of the percentage of bleb reabsorption events for UNT, serum starved, serum add back, VPA, VPA serum starved, and VPA serum add back MEF cells. (D) Example images and graph of bleb-based (dark green) vs. non-bleb based (light green) nuclear rupture UNT, serum starved, serum add back, VPA, VPA serum starved, and VPA serum add back in MEF cells. (E) Example images of MEF NLS-GFP time lapse imaging of a nuclear rupture occurring three times in the same nucleus. (F, G) Graphs of (F) percentage of nuclei the rupture and (G) nuclear rupture frequency denoting the number of times a single nucleus rupture over the 3-hour timelapse for wild type untreated (UNT), serum starved, and serum add back as well as VPA, VPA serum starved, and VPA serum add back in MEF cells. N=3 biological replicates with >100 nuclei for all graphs. Error bars represent standard error and statistical tests are one-way ANOVA with a post-hoc Tukey test except panel A within conditions (10% to 0.5%) used a two-tailed paired Student’s t-test, with significance denoted by *= p<0.05, **= p<0.01, and ***=p<0.001. Scale bar =10µm.

We measured nuclear blebbing percentages to determine if the decreased transcriptional activity in serum starved conditions suppressed nuclear blebbing. Wild type untreated MEFs (UNT) exhibited a blebbing percentage of approximately 4.4 ± 0.5% (**Figure 2A**). Serum starvation for 72 hours significantly decreased the percentage of nuclear blebbing to 3% ± 0.1 (**Figure 2A**). Chromatin and lamin perturbations are known to result in higher blebbing percentages (Lammerding *et al*., 2006; Vargas *et al*., 2012; Hatch and Hetzer, 2016; Stephens *et al*., 2019b; Pho *et al*., 2023). Chromatin decompaction via histone deacetylase inhibitor VPA and lamin perturbation lamin B1 KO (*Lmnb1-/-*) bleb significantly more than UNT, about 10%.

Serum starvation for 72 hours of VPA-treated and *Lmnb1-/-* cells showed a significant decrease in blebbing percentage to 4%, similar to untreated wild type (**Figure 2A**). Thus, decreased transcriptional activity via serum starvation significantly decreased nuclear blebbing in both wild type and nuclear perturbations.

Transcription activity and nuclear blebbing is restored within hours upon serum add back to serum starved cells. Upon serum add back (10% FBS) MEF wild type untreated, MEF VPA treated, and MEF *Lmnb1-/-* cells were time lapsed at 3-minute intervals and nuclear blebbing percentages were reported at 30-minute intervals. Over the course of a 3-hour time lapse, we saw blebbing percentages increase significantly for each condition (**Figure 2B**). Nuclear blebbing percentages were returned to similar levels as untreated in 2 hours. Like the MEF cell lines, HT1080 cells exhibited a decrease in nuclear blebbing following serum starvation and an increase in nuclear blebbing after serum added back (**Supplemental Figure 2, A and B**). In summary, decreased transcriptional activity via serum starvation decreases nuclear blebbing, while adding serum back restored transcriptional activity and nuclear blebbing.

To assess whether serum manipulation influenced nuclear bleb stability, we quantified the persistence of nuclear blebs throughout interphase. We defined bleb stability as the ability of a nuclear bleb to persist until the end of the 3-hour time lapse or mitosis. Blebs that disappeared were categorized as reabsorbed and blebs that persisted throughout the time lapse were classified as stable. In UNT MEFs, nuclear blebs were highly stable, with only about 10% reabsorbing (**Figure 2C**). Serum starvation resulted in a substantial increase in bleb reabsorption, with approximately 75% of nuclear blebs being reabsorbed (**Figure 2C**). Following serum add back, bleb reabsorption decreased significantly to approximately 25% (**Figure 2C**). These results indicate that transcription activity modulated via serum controls nuclear bleb stability.

In addition to changes in nuclear blebbing, nuclear rupture dynamics were examined. Nuclear ruptures were defined as a greater than 25% increase in the cytoplasm/nucleus ratio of nuclear localization signal green fluorescence protein (NLS-GFP) (Pho *et al*., 2023). Nuclear ruptures can occur due to nuclear blebs, denoted as bleb-based, or in normally shaped nuclei, denoted as non-bleb-based. In UNT MEF cells approximately 75% of total ruptures are bleb-based.

Serum-starved cells significantly decreased to 35% bleb-based ruptures (**Figure 2D**). Serum add back restored the number of bleb-based ruptures to UNT levels within three hours. In VPA-treated cells, a similar outcome occurred as bleb-based ruptures were dependent on transcription activity (**Figure 2D**). Overall, transcriptional activity alters the type of nuclear rupture.

Nuclear ruptures cause dysfunction (Stephens, 2020). To determine how serum manipulation-based changes in transcription activity influence nuclear ruptures, we measured both the percentage of nuclei that rupture and then how many times a given nucleus ruptures over the 3-hour timelapse (**Figure 2E**). As previously established, we measured an increase in the percentage of nuclei that rupture as well as how frequently a nuclear rupture occurs from UNT to VPA. Untreated wild type 4% of the nuclei rupture with a frequency of about 1.5 which significantly increased in VPA-treated to 9% of nuclei rupture and 2.5 rupture frequency for a given nucleus (**Figure 2, F and G**). In MEF untreated serum starvation and add back did not change the percentage of nuclei of rupture but did alter frequency, which was decreased by starvation then restored by add back (**Figure 2F**). In MEF VPA-treated nuclei serum starvation decreased nuclear rupture percentages and frequency to wild type levels, while 3-hour serum add back was insufficient to restore both rupture percentage and frequency (**Figure 2G**). Thus, perturbations that increase nuclear ruptures and their frequency are dependent on transcriptional activity.

### Active RNA Polymerase II is enriched in nuclear blebs

To assess whether transcriptional activity is a key component of nuclear deformations, we quantified the levels of active RNA Pol II in the nuclear bleb. To determine possible nuclear bleb enrichment, we compared the levels of Hoechst DNA stain signal and active RNA Pol II markers pSer5 and pSer2 in the nuclear bleb versus the nuclear body. In all conditions, as expected, we found that the DNA density was decreased in the nuclear bleb relative to the body (**Figure 3, E and F**), as previously reported (Stephens *et al*., 2018; Bunner *et al*., 2024; Chu *et al*., 2025). For transcriptional initiation marker RNA Pol II pSer5, we found that it was enriched relative to DNA in all nuclear blebs, regardless of perturbation (**Figure 3F**). We found that transcriptional elongation marker RNA Pol II pSer2 varied in the enrichment of the nuclear bleb depending on the perturbation (**Figure 3F**). These data agree with our previous findings (Berg *et al*., 2023). Overall, enrichment in the nuclear bleb suggests that transcriptional activity is a core component of nuclear blebs as it is important for both bleb formation and stability.

**Figure 3.**
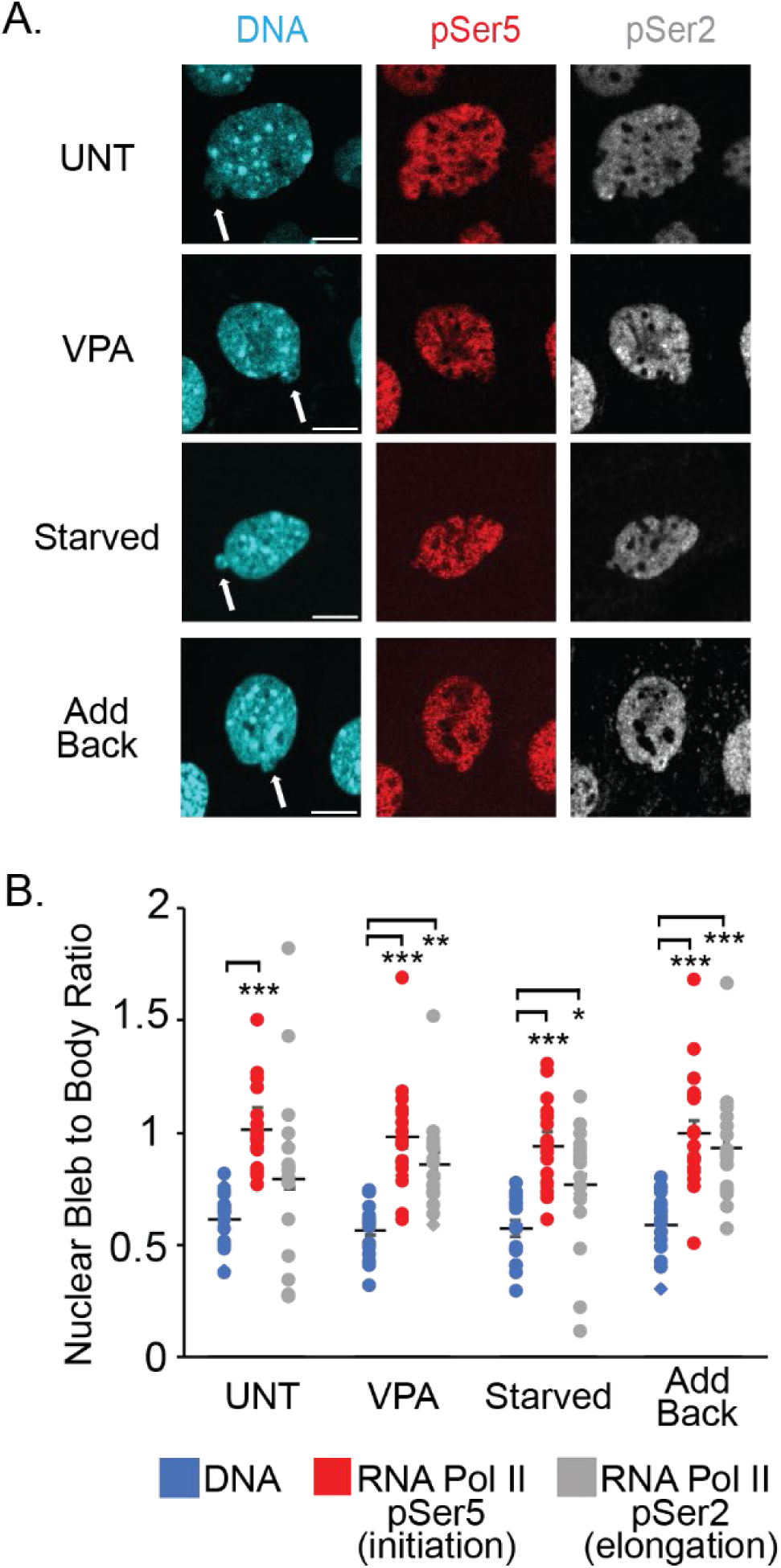
Nuclear bleb vs. body intensity measurements of active RNA Pol II. (A) Example images of DNA via Hoechst and active RNA Pol II markers pSer5 (initiation) and pSer2 (elongation) in MEF WT cells UNT, VPA, serum starved or add back conditions. Images contrasted for visibility, not reflective of relative intensity changes across conditions. (B) Graph of bleb to body relative fluorescence ratios for DNA and active RNA Pol II pSer5 initiation and pSer2 elongation. N=3 biological replicates with >18 nuclei for all graphs. Error bars represent standard error and statistical tests are one-way ANOVA with a post-hoc Tukey test with significance denoted by *= p<0.05, **= p<0.01, and ***=p<0.001. Scale bar =10µm.

### Serum modulated transcriptional activity does not change actin confinement or contraction

To determine the mechanism by which transcriptional activity modulated by serum affects nuclear blebbing, we assessed its impact on actin. Actin confinement and contraction are known to antagonism nuclear shape and contribute to nuclear blebbing (Khatau *et al*., 2009; Hatch and Hetzer, 2016; Mistriotis *et al*., 2019; Pho *et al*., 2023). Nuclear height is measured by spinning disk confocal microscopy of DNA stained by Hoechst to provide an indirect measure of actin confinement, with smaller nuclear heights indicating increased confinement. Untreated MEF wild type cells measured an average nuclear height of 4.4 ± 0.3 µm (**Figure 4A**). Nuclear height did not significantly change following serum starvation or add back compared to wild type conditions.

**Figure 4.**
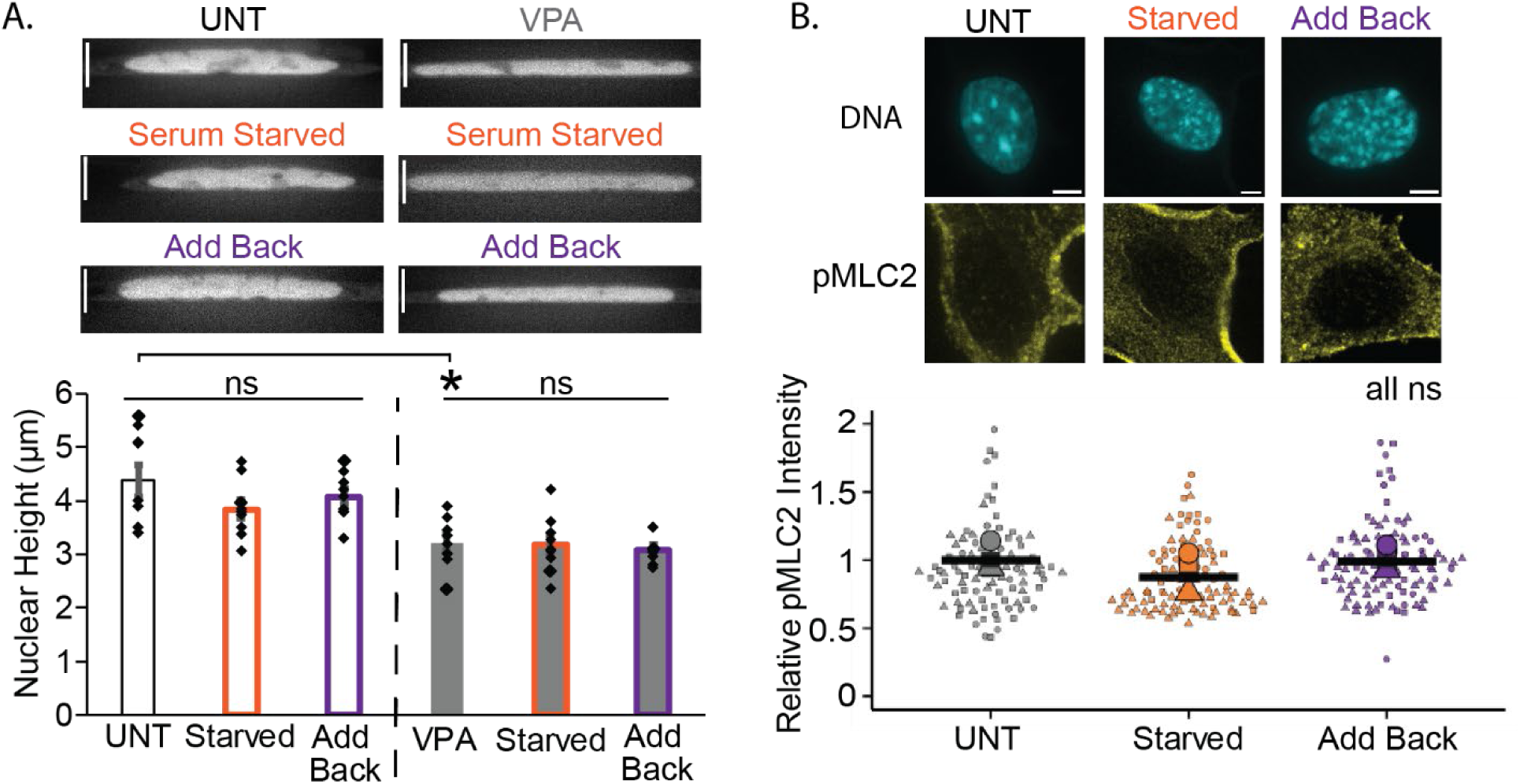
Serum manipulation does not affect nuclear height or pMLC2 levels. (A) Example images of nuclei-stained DNA via Hoechst and graph of nuclear height (µm) for wild type untreated (UNT), serum starved, and serum add back as well as VPA, VPA serum starved, and VPA serum add back in MEF cells. n = 20 nuclei measured for each condition. (B) Example images and graph of relative fluorescence intensity of pMLC2 immunofluorescence staining in UNT, serum starved, and serum add back MEF cells. N=3 biological replicates with >20 nuclei for all graphs. Error bars represent standard error and statistical tests are one-way ANOVA with a post-hoc Tukey test, with significance denoted by *= p<0.05, **= p<0.01, and ***=p<0.001. Scale bar =10µm.

The histone deacetylase inhibitor valproic acid (VPA) results in a weaker nucleus less capable of resisting actin confinement, resulting in reduced nuclear height (Berg *et al*., 2023; Pho *et al*., 2023). In MEF cells treated with VPA, nuclear height was significantly decreased to 3.2 ± 0.2 µm relative to untreated cells (**Figure 4A**), recapitulating previous publication results and confirming our ability to measure changes in nuclear height. Serum modulation of VPA-treated MEF cells measured no significant change in nuclear height (**Figure 4A**). Overall, this data suggests that actin confinement is not responsible for the changes in nuclear blebbing seen in serum starvation and add back.

Another actin antagonizing force is actin contraction that can be quantified through immunofluorescence of phosphorylated myosin light chain 2 (pMLC2, (Murrell *et al*., 2015)). Relative pMLC2 intensity measurements showed no significant changes in actin contraction levels between untreated, serum-starved, and serum add back cells (**Figure 4B**). Furthermore, using fluorescent ubiquitination-based cell cycle indicator (FUCCI) cells or Cdt1 immunofluorescence, we measure no change in cell cycle distribution (**Supplemental Figure 3**), which can lead to changes in nuclear blebbing (Bunner *et al.,* 2025 bioRxiv 2025.04.16.649171). This data suggests that actin contraction and cell cycle are not contributing to the changes in nuclear blebbing that occur upon serum starvation and add back.

### Transcriptional activity modulation does not change chromatin or lamin based nuclear mechanical resistance

Chromatin and lamin alterations are major contributors to nuclear blebbing through their role providing nuclear stiffness (Stephens *et al*., 2019a). Increased levels of euchromatin via HDACi VPA have been shown to promote nuclear blebbing by weakening chromatin-based nuclear stiffness (Stephens *et al*., 2018; Hobson *et al*., 2020; Kalinin *et al*., 2021). To assess whether chromatin histone modifications were altered by serum manipulation, immunofluorescence staining for the euchromatin marker H3K9ac was performed. Untreated MEF cells exhibited no significant changes in H3K9ac levels following serum starvation and add back compared to untreated cells (**Figure 5A**). VPA-treated cells showed a significant increase in euchromatin levels compared to untreated, as expected. Similar to untreated, VPA-treated cells that were serum starved and added back did not significantly change (**Figure 5A**). Thus, serum modulation does not affect euchromatin H3K9ac levels to alter levels of nuclear blebbing.

**Figure 5.**
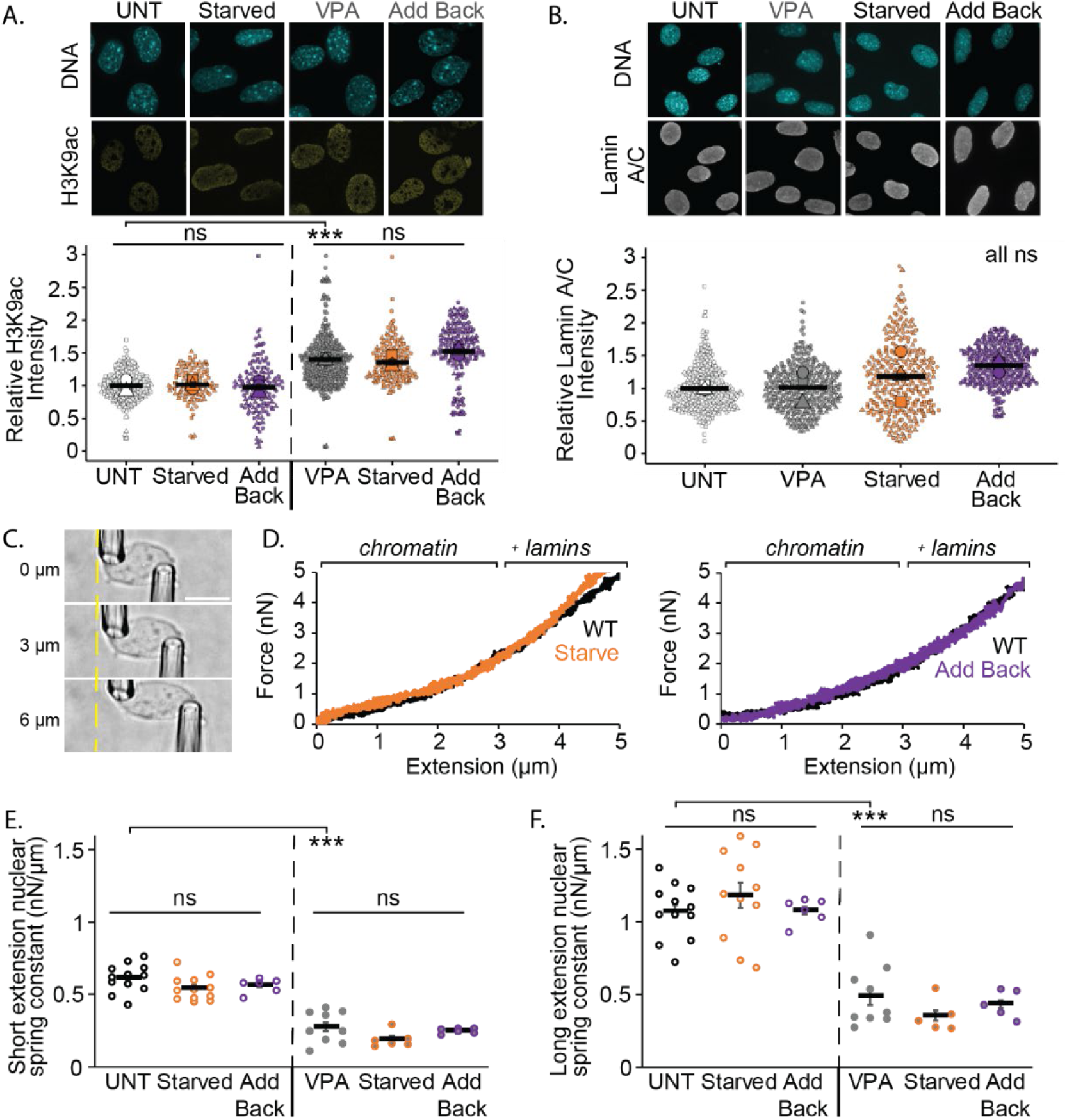
Serum manipulation does not change euchromatin levels, lamin levels, or the nuclear spring constant. (A) Example images of nuclei stained with Hoechst for DNA and H3K9ac euchromatin marker. Graph of relative H3K9ac intensities for MEF wild type untreated (UNT), serum starved, and add back as well as VPA, VPA starved, and VPA serum add back. (B) Example images of nuclei stained with Hoechst for DNA and lamin A/C and graph of relative lamin A/C intensities UNT, VPA, serum starved, and serum add back MEF WT cells. N=3 biological replicates with >20 nuclei. (C) Example images of isolated MEF vimentin null nucleus dual micropipette micromanipulation force-extension experiment at 0, 3, and 6 µm nucleus extension. The right pull pipette extends (µm) the nucleus as the left force pipette’s deflection multiplied by its premeasured bending constant measures force (nN). (D) Example nucleus force-extension curves where brackets denote the 0-3 μm short extension (chromatin) and > 3 μm long extension (chromatin + lamins) regimes. Average nuclear spring constant (µm/nN) for (E) short extension <3 µm and (F) long extension >3 µm for untreated UNT and VPA-treated at serum normal, starved, and add back (n = 12,12,6, 9, 6, 6 nuclei respectively). Error bars represent standard error and statistical tests are one-way ANOVA with a post-hoc Tukey test, with significance denoted by *= p<0.05, **= p<0.01, and ***=p<0.001. Scale bar =10µm.

Lamin A/C plays a crucial role in nuclear mechanics and rigidity, which can influence nuclear blebbing (Vahabikashi *et al*., 2022; Pho *et al*., 2024). Lamin A/C levels were measured using immunofluorescence to determine the possible changes upon serum modulation. No significant differences in lamin A/C levels were observed between untreated, VPA, serum-starved, or serum add back (**Figure 5B**). Overall, this data suggests that changes in lamin A/C levels are not responsible for serum modulation nuclear blebbing effects.

Dual micropipette micromanipulation nucleus force measurements were conducted to further assess potential changes in chromatin and lamin contributions to nuclear stiffness. These measurements allow for the separation of the relative roles of chromatin and lamins in nuclear mechanical strength. For micromanipulation experiments, MEF vimentin-null (MEF V-/-) cells were used because they allow easy nucleus isolation (Currey *et al*., 2022). The isolated nucleus is suspended between micropipettes. The nucleus is extended by one micropipette while force is measured by the deflection of the other micropipette multiplied by a pre-measured bending constant (**Figure 5C**).

Dual micropipette micromanipulation nucleus force measurements reveal no changes upon serum manipulation. Serum starvation did not result in significant changes in either the chromatin dominated short extension regime or the long extension strain stiffening lamin regime compared to untreated wild type (**Figure 5D-F**). Serum add back nuclear spring constant for both short and long regimes were similar to both wild type untreated and serum-starved nuclei. VPA-treated cells exhibited a significant decrease in nuclear stiffness relative to untreated wild type, consistent with previous findings (Shimamoto *et al*., 2017; Stephens *et al*., 2017; Hobson *et al*., 2020; Currey *et al*., 2022). Both serum starvation and add back treated with VPA also showed no significant differences from VPA-treated MEF V-/-cells (**Figure 5, E and F**). Taken together both immunofluorescence levels and micromanipulation nucleus force measurements upon serum manipulation reveal no changes in chromatin and lamin A/C, the two key resistive elements of the nucleus.

### Transcriptional activity regulates chromatin motion

We previously hypothesized that transcriptional activity controls chromatin motion which leads to changes in nuclear blebbing supported by a physics-based simulation model (Berg *et al*., 2023). However, a direct measure of transcription activity’s effect on individual chromatin domain motion via mean squared displacement remains unknown. To measure chromatin motion, we transfected MEF cells with fluorescent Cy3-dUTPs that labeled chromatin replication domains and then tracked foci movement via mean squared displacement.

To measure levels of chromatin motion we live cell imaged transfected Cy3-dUTPs at 2-second intervals for 150 seconds and used the TrackMate software in ImageJ (Tinevez *et al*., 2017; Ershov *et al*., 2022). To account for changes in nuclear movement, we cropped individual nuclei and performed image registration through the NLS-GFP channel in FIJI to remove nuclear body movement. Once the nuclei were registered, they were analyzed via the TrackMate plugin to track chromatin domain movement. To qualitatively see the change in chromatin motion we graph an example tracks of single chromatin domain’s motion on the order of 1 µm (**Figure 6A**). Then we analyzed the chromatin motion data quantitatively using mean squared displacement (MSD). Cy3-dUTP-labeled chromatin domains in MEF wild type untreated measure MSD of 0.12 µm^2^ ± 0.01 over a seventy-five-second interval (**Figure 6B**). This is our control condition to see if serum modulations or other drugs can change the chromatin motion.

**Figure 6.**
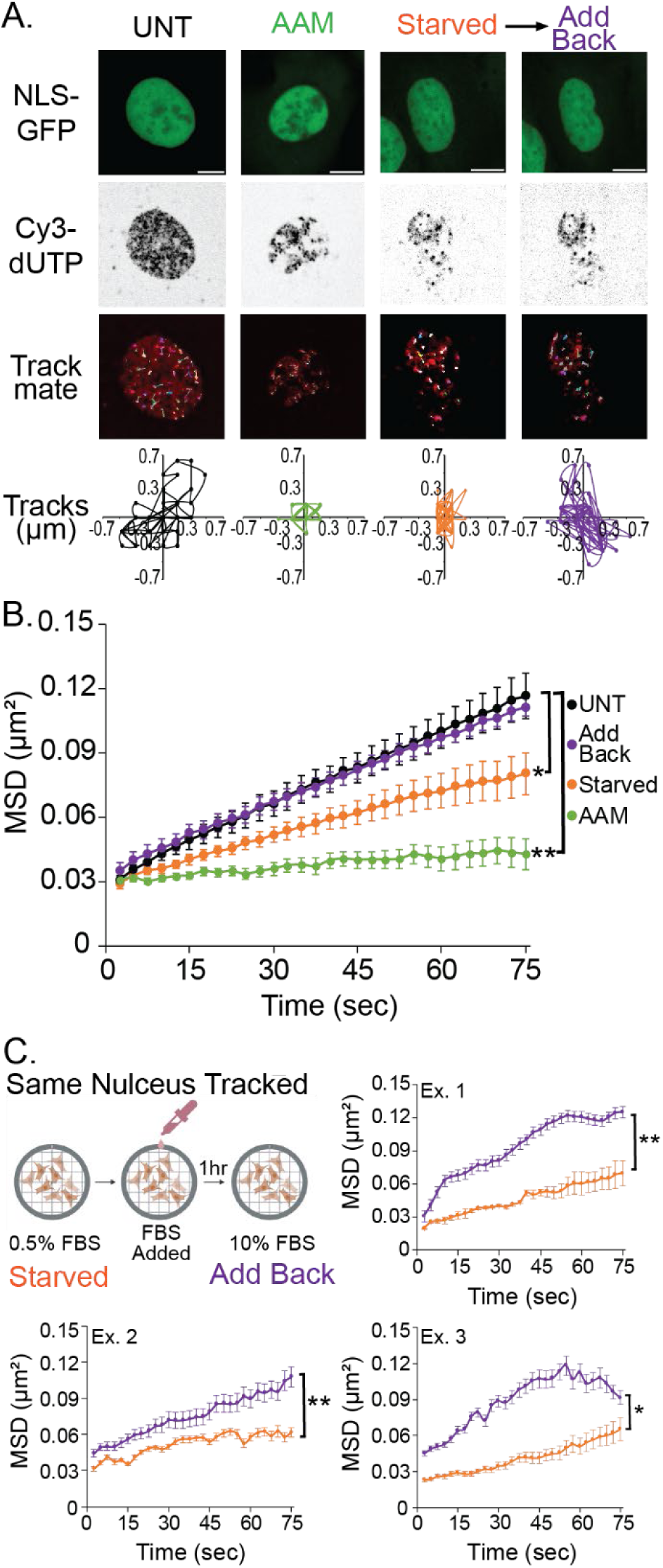
Chromatin motion is decreased by serum starvation and restored by serum add back. (A) Example images of wild type MEF nuclei with NLS-GFP and Cy3-dUTP used for tracking chromatin motion. Example Trackmate tracks for all measured chromatin foci domain motion tracked and example single chromatin domain foci track for untreated control (UNT, black), RNA Pol II inhibition (AAM, green), serum starved (orange), and serum add back (purple). (B) Mean squared displacement (MSD) plots of UNT, AAM, serum starved, and serum add back of >8 foci averaged per nucleus over 75 seconds, which is half of full tracking of 150 seconds, with 20 nuclei per condition, except AAM with 10 nuclei. (C) Three example mean squared displacement (MSD) plots of the same nucleus serum starved (orange) and serum added back (purple) of >8 foci averaged per nucleus over 75 seconds, which is half of full tracking of 150 seconds, for 3 nuclei. Serum starved and serum add back are paired measurements of the same nucleus before and after serum re-addition. Error bars represent standard error and statistical tests are one-way ANOVA with a post-hoc Tukey test except panel C Two-tailed paired Student’s T-test, with significance denoted by *= p<0.05, **= p<0.01, and ***=p<0.001. Scale bar =10µm.

First, we determined the bulk role of transcriptional activity in chromatin motion via the RNA Pol II inhibitor alpha amanitin (AAM). We found that AAM-treated cells exhibited significantly decreased chromatin domain MSD when compared to untreated wild type (**Figure 6, A and B**). This data agrees nicely with published studies that transcription inhibition decreases coherent chromatin motion (Shaban *et al*., 2018, 2020). Thus, transcription inhibition drastically decreases chromatin domain motion via MSD.

Next, we established the role of transcription activity modulation through serum starvation and add back. To better control for changes from serum starvation to add back, we imaged the same nuclei of cells serum starved for 72 hours before and after serum add back for 1 hour, by using a gridded single well imaging dish (**Figure 6C**). Serum starved cell nuclei had significantly decreased chromatin domain MSD when compared to untreated wild type cells (**Figure 6, A and B**). These cells with serum added back for one hour increased chromatin domain MSD motion relative to serum starved, returning MSD back to untreated wild type levels. The restoration of chromatin domain MSD motion of the same nucleus was consistently significant from serum starved and then add back across individual nuclei (**Figure 6C**). This can be recapitulated in a second cell type as wild type HT1080 cells also showed a significant increase in chromatin motion when comparing the same cell serum starved to serum add back after one hour (**Supplemental Figure 2, C and D**). Thus, serum modulation of transcriptional activity regulates chromatin motion providing a mechanism by which to drive nuclear blebbing.

### BO2 increased chromatin motion and nuclear blebbing independent of nuclear stiffness and transcriptional activity

Nuclear blebbing could be driven by transcriptional activity through chromatin motion or through an alternative process. To determine that chromatin motion controls nuclear blebbing, we would need a scenario in which chromatin motion increased independent of transcription. It has been established that chromatin motion increases upon treatment of cells with BO2 a RAD51 inhibitor (Maarouf *et al*., 2024). To verify these findings, we treated Cy3-dUTP transfected cells with BO2 10 µM for 24 hours and track chromatin domain MSD. BO2-treated nuclei exhibited a significantly increased MSD relative to untreated controls cells (**Figure 7, A and B**). Measurements of active RNA Pol II pSer5 and pSer2 were not altered by B02-treatement (**Supplemental Figure 1**). Thus, BO2 increases chromatin domain motion measured by MSD without altering transcriptional activity.

**Figure 7.**
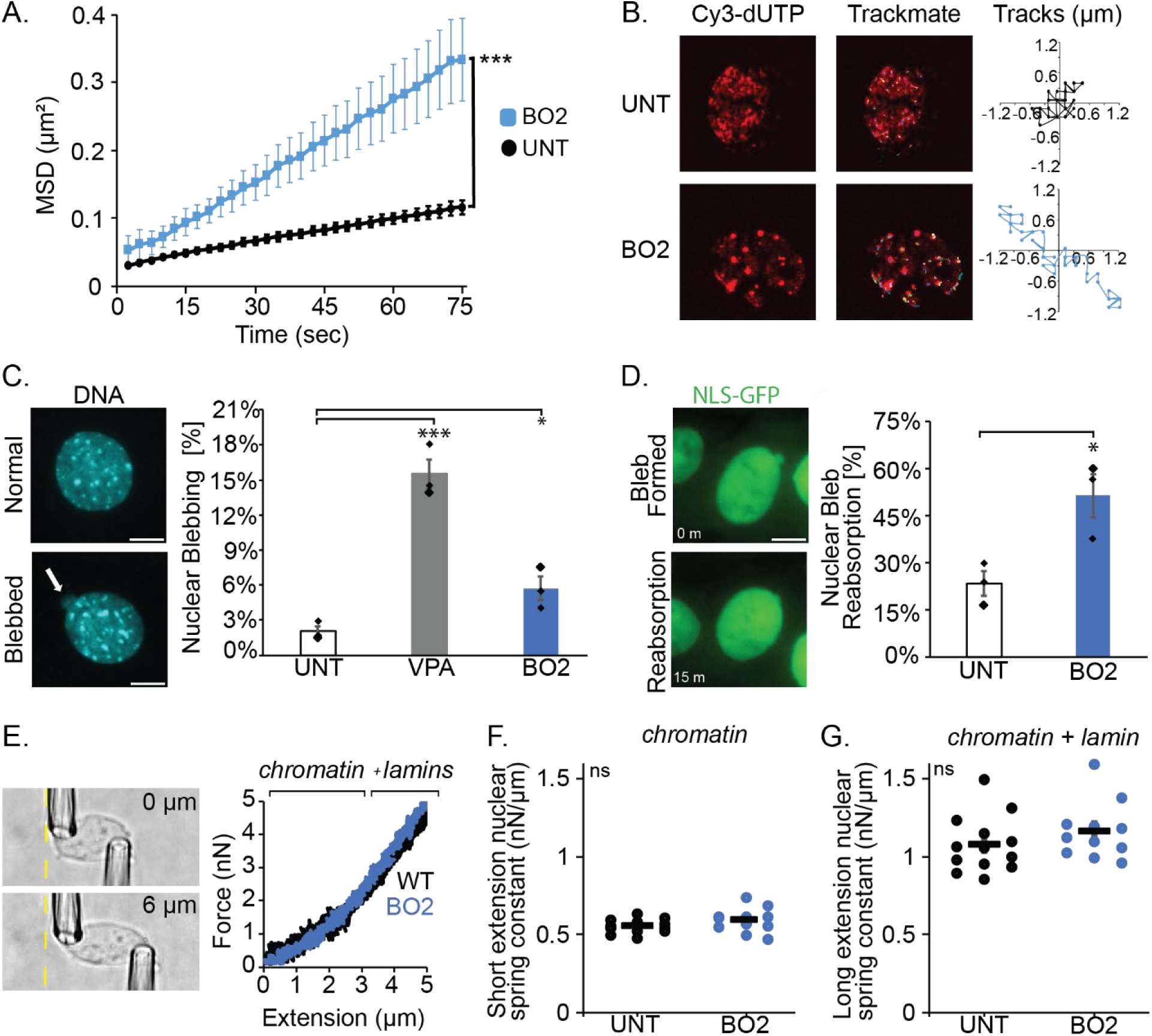
B02 increases chromatin motion and nuclear blebbing but not the nuclear spring constant. (A) Mean squared displacement (MSD) plots of UNT, and BO2 of >8 foci averaged per nucleus over 75 seconds, which is half of full tracking of 150 seconds, 10 nuclei per condition. (B) Example images of wild type MEF nuclei with CY3-dUTP used for tracking chromatin motion. Example trackmate tracks for all measured chromatin foci domain motion tracked and example single chromatin domain foci track for untreated control (UNT, black) and BO2 (blue). (C) Example images of normal and blebbed MEF wild type nuclei stained with Hoechst and graph of blebbing percentage in normal (black outline), VPA (gray fill), and BO2 (blue fill). (D) Example images of a MEF nuclear bleb reabsorption and graph of the percentage of bleb reabsorption events for UNT and BO2 treated MEF cells. For C and D N=3 with n> 50 cells precondition. (E-G) Dual micropipette micromanipulation nuclear spring constant measurements for (X) short extension > 3 µm and (X) long extension < 3 µm for MEF untreated (UNT, n = 13) and B02-treated (n = 11) nuclei. Error bars represent standard error and statistical tests are two-tailed (C,D) paired or (A, F, G) unpaired Student’s t-test with significance denoted by *= p<0.05, **= p<0.01, and ***=p<0.001. Scale bar =10µm.

The hypothesis that chromatin motion is essential to nuclear bleb formation suggests that increased chromatin motion via B02 should increase nuclear blebbing. Untreated and cells treated with BO2 for 24 hours were imaged via DNA stain Hoechst to determine the percentage of nuclear blebbing. Relative to untreated controls, BO2-treatment resulted in increased nuclear blebbing from 2.0 % to 5.7 % (**Figure 7C**). To elucidate the stability of these blebs, we time lapsed imaged untreated and BO2-treated cells for 3 hours via NLS-GFP. Nuclear bleb reabsorption increased in BO2-treated cells compared to untreated (**Figure 7D**). Thus, the data show that increased chromatin motion is sufficient to increase nuclear blebbing, although those blebs are unstable.

To ensure that RAD 51 inhibition via BO2 induced nuclear blebbing is not due to changes in the nuclear mechanics, we measured the nuclear spring constant. Micromanipulation force measures revealed that untreated and BO2-treated nuclei had similar nuclear rigidity (**Figure 7E-G**). Specifically, the nuclear spring constant remained unchanged for both the short extension dominated chromatin regime and long extension regime with chromatin plus lamin A-based strain stiffening (**Figure 7, F and G**). Immunofluorescence of eu-/heterochromatin and lamin levels in BO2 treated cells did not significantly change relative to untreated wild type (**Sup Figure 1 B-C**). Thus, BO2 increased chromatin motion is a mechanism that can drive nuclear blebbing independent of nuclear stiffness and transcriptional activity.

## Discussion

We detail that transcription-activity-generated chromatin motion is a new contributor to nuclear blebbing through modulation of transcriptional activity via serum manipulation. This finding provides a major update to our understanding of nuclear blebbing, which is not solely caused by an imbalance in nuclear stiffness resisting actin confinement and contraction. Furthermore, these novel findings inform our understanding of how a nuclear bleb forms via chromatin domain level motion. These findings are important for understanding core fundamental cell biology and human disease. Many human diseases present both increased transcriptional activity and abnormal nuclear morphology. We reveal that increased transcriptional activity can result in nuclear blebbing, known to drive nuclear dysfunction that likely advances the disease state.

### Transcription regulates chromatin domain motion to form nuclear blebs

Transcription activity impacts chromatin motion differently on nano vs micro scales. Transcription activity decreases single nucleosome movement (Nagashima *et al*., 2019). We find the opposite for chromatin replication domains labeled via Cy3-dNTPs (**Figure 6**). This agrees with transcriptional activity increasing micron-scale coherent chromatin motion (Shaban *et al*., 2018), the chromatin fractional moving mass via partial wave spectroscopy (Almassalha *et al*., 2025) and motion of both promotors and enhancers (Gu *et al*., 2018). RAD51 inhibitor BO2 treatment alters chromatin domain motion and not nucleosome motion (Maarouf *et al*., 2024). Thus, our data shows that increasing chromatin domain motion is the essential behavior behind nuclear blebbing (**Figure 7**).

The scale and chromatin linking agree with RNA polymerase II-generated chromatin domain motion causing nuclear blebbing. Nuclear blebs are defined as >1 µm sized protrusions which is on the order of chromatin domain and coherent chromatin motion 1-3 µm (Zidovska *et al*., 2013; Shaban *et al*., 2018, 2020; Locatelli *et al*., 2022; Chu *et al*., 2024). Nuclear blebs also form and grow on the minute timescale (Berg *et al*., 2023). Chromatin is highly linked via specific proteins such as HP1α (Strom *et al*., 2021; Williams *et al*., 2024) and Hi-C type chromatin interactions about every 20 kb (Belaghzal *et al*., 2021). Simulations show that modeling chromatin crosslinkers as slow, or long lived, increases domain motion, while fast, short lived, crosslinkers suppress motion (Maarouf *et al*., 2024). Transcriptional proteins, including RNA Pol II and mRNA have been reported to make chromatin crosslink-like interactions. Though our data shows that changes in transcriptional activity do not affect nuclear spring constant, suggesting no significant change in chromatin crosslinking due to transcriptional activity (**Figure 5**). Thus, transcription activity generated chromatin domain motion and shown chromatin linking aids the scale of chromatin movement required to form and stabilize a nuclear bleb.

Physics-based simulation modeling supports the experimental findings. A mechanical model of the nucleus consisting of chromatin, lamins, chromatin-chromatin linkers, and chromatin-lamin linkers was first generated (Banigan *et al*., 2017) and then further developed (Strom *et al*., 2021). Addition of chromatin interacting motors was then shown to increase nuclear shape fluctuations (Liu *et al*., 2021) and bleb-like bulges (Berg *et al*., 2023). Joint, or coherent, chromatin movement pushing out the nuclear periphery underlies the formation of these bulges. These physical modeling outcomes are recapitulated by our experiments. Modeling of the number of motors is equivalent to number of active motors experimentally measured (**Figure 1**) which cause proportional changes in nuclear blebbing (**Figure 2**) through the mechanism of chromatin motion (**Figure 6 and 7**).

### Transcription activity-based chromatin motion as a new contributor to nuclear blebbing

Previously, our understanding of how a nuclear bleb forms was due to an imbalance of nuclear stiffness and actin antagonism. This model fails to clarify how a nuclear bleb forms. One major theory is that actin confinement could rupture the nucleus allowing chromatin to spew into a herniation. However, this theory has no supporting data and only data refuting it. The majority of nuclear blebs form first and then this high curvature causes rupture as a secondary step. This statement is supported also by the fact that laser induce nuclear ruptures do not form nuclear blebs (Halfmann *et al*., 2019). Confinement via artificial confinement devices rupture nuclei but do not generate nuclear bleb-like structures. Instead, artificial confinement forms thin and length membrane-based protrusions that rupture but are reabsorbed quickly, unlike nuclear blebs that remain for hours if not the whole cell cycle. Thus, nuclear resistance and actin antagonism is insufficient to explain nuclear bleb formation.

Understanding that transcription driven chromatin motion underlies nuclear bleb formation provides a nucleation force. Specifically, chromatin colliding with the nuclear periphery can push out to provide a herniation. The composition of a nuclear bleb agrees with transcription as a new contributor. First, nuclear blebs are best defined by decreased DNA density (Bunner *et al*., 2024; Pujadas Liwag *et al*., 2025) and increased euchromatin (Stephens *et al*., 2018) which both are associated with transcription activity. Second, we provide data that nuclear blebs are enriched in transcription activity marked by active RNA Pol II pSer5/2 (**Figure 3**), supported by previous publications (Bercht Pfleghaar *et al*., 2015; Berg *et al*., 2023). More transcriptionally active chromosomes are more likely to end up in bleb (Bercht Pfleghaar *et al*., 2015). In prostate cancer cells LNCaP treatment of testosterone (DHT) increased nuclear blebbing where active androgen responsive genes are enriched in the nucleus bleb (Helfand *et al*., 2012). Overall, the data support that transcription is an essential component of nuclear blebbing and now we have provided the mechanism of chromatin motion underlying this behavior.

### Nuclear shape is determined by force balance and chromatin motion

Transcription activity driven chromatin motion and nuclear vs. actin force balance are equally important factors for nuclear blebbing and rupture (**Figure 8**). This can best be seen in the case of BO2 treatment. Increase of chromatin motion via B02 can increases nuclear blebbing but only marginally and the blebs are unstable (**Figure 7**). The reverse can example is VPA-treatment in which the nucleus is weaker disrupting the nuclear-actin force balance, but transcriptional activity motion is required to form and stabilize nuclear blebs (**Figure 2 and 5**). Thus, the data support that both chromatin motion and a disrupted nuclear-actin force balance are pivotal to nuclear blebbing and stability.

**Figure 8.**
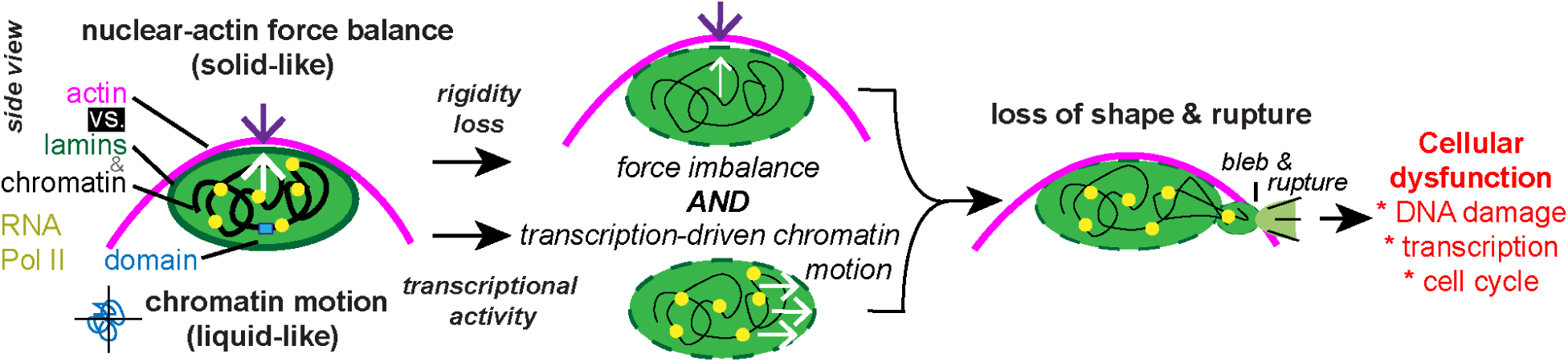
Nuclear rigidity and chromatin motion are the two key contributors to nuclear shape maintenance. Side view schematic of the nucleus. Nuclear mechanical components chromatin and lamins resists actin and external antagonism to maintain shape (top). We reveal a new contributor that affects nuclear shape via transcriptional activity generated chromatin motion (bottom). When the nucleus is weaker via chromatin or lamin perturbation and there is sufficient transcriptional activity produced chromatin motion the nucleus will form a nuclear bleb and rupture known to cause dysfunction. These two contributors to nuclear shape represent the solid-like nuclear properties acting as a spring to resists and liquid-like properties of motion and flow. Bottom right inset: RNA Pol II is a chromatin motor protein capable of generating forces on the chromatin to generate movement, aided by known chromatin linking, that regulates chromatin domain motion to push against and deform the nuclear periphery to make a nuclear bleb.

Chromatin has varying physical properties based on size scale. Nuclear force measurements clearly show that chromatin on the genome scale is a major mechanical component (Stephens *et al*., 2017; Hobson *et al*., 2020; Nava *et al*., 2020; Currey *et al*., 2022; Bergamaschi *et al*., 2024; Williams *et al*., 2024). Since chromatin on the nuclear scale has a bulk modulus, it acts as a solid. However, force measurements of single chromatin loci show a liquid-like behavior (Keizer *et al*., 2022). Photobleaching of heterochromatin and euchromatin show solid-like behaviors where chromatin proteins show liquid-like behavior (Strickfaden *et al*., 2020). In agreement, analysis of single nucleosome movement up to replication domain foci also show liquid-like behavior where a 6 µm diameter region shows solid-like behavior (Nozaki *et al*., 2023). Our data verify these previous findings through chromatin’s solid-like capacity to provide a nuclear spring constant (**Figure 5 and 7**) that is separable from chromatin’s liquid-like capacity to flow via chromatin motion (**Figure 6 and 7**). Thus, chromatin is a gel with solid-like restive properties on the macro scale needed to resist nuclear shape deformation and a liquid-like micro (< 1-3 µm) and nano scale that can flow to allow nuclear bleb formation (**Figure 8**).

### Implications for transcriptional activity diving nuclear blebbing in disease

Many human diseases are associated with abnormal nuclear morphology and increased transcription activity. Specifically, cancers are well-known to have increased transcription via transcription factors or hormones. For example, c-Myc is a transcription factor largely associated with cancer (Dhanasekaran *et al*., 2022). Less broad and more specific, testosterone (DHT) is well known to active the androgen receptor which then activates androgen responsive genes (Tan *et al*., 2015). This activation of a specific transcription response is sufficient to increase nuclear blebbing in LNCaP prostate cancer cells (Helfand *et al*., 2012). Increased nuclear blebbing correlates with Gleason score which is used to determine cancer aggressiveness. Other transcription factors are implicated in cancer and are current targets of therapies (Bushweller, 2019). Thus, transcription activation in these diseases could provide the nucleating force to cause nuclear deformation and ruptures that cause nuclear dysfunction and drive disease state alongside their functions in determining gene expression.

We provide that modulation of transcription activity globally affects chromatin motion to drive nuclear blebbing. Future work could further address the capacity of a single transcription factor or hormone response to generate sufficient changes in chromatin motion and nuclear blebbing. Another goal of future studies will be to address the thresholds of the amount of chromatin undergoing motion and amount of motion required to cause nuclear blebbing and rupture.

Finally, future work will be required to understand the event of nuclear bleb formation given that we have now provided chromatin motion as a key driving force. This work advances our understanding of the basic biology behind nuclear deformations that are a key hallmark and contributor to cellular dysfunction in human diseases.

## Materials and Methods

### Cell culture

Mouse Embryonic Fibroblast (MEF) were previously described in (Shimi *et al*., 2008; Stephens *et al*., 2018; Vahabikashi *et al*., 2022). MEF wild-type (WT and lamin B1 knockout (LMNB1-/-) cells were cultured in DMEM (Corning) completed with 10% fetal bovine serum (FBS, HyClone) and 1% penicillin/streptomycin (Corning). Cells were grown and incubated at 37 C and 5 % CO2, passaged every 2 to 3 days. Human fibrosarcoma HT1080 cells were cultured and passaged similarly. HT1080 cells were obtained from Orth lab and had a Fluorescent Ubiquitin Cell Cycle Indicator (FUCCI) with lentiviral plasmids mKO2-hCdt1(30/120) [DDBJ/EMBL/GenBank, AB370332] in pCSII-EF vector and mAG-hGem(1/110) [DDBJ/EMBL/GenBank, AB370333] in pCSII-EF.

For serum starvation, cells were grown in imaging dishes with cell culture media made with 0.5% FBS. Cells were starved for 48-72 hours. Serum stimulation was done by adding FBS back to imaging dishes to get back to 10% FBS. All cell lines were tested for contamination weekly and were obtained from ATCC or authenticated before beginning experiments.

### Biochemical treatments

MEF WT cells were treated with either 4 mM valproic acid (VPA, 1069-66-5, Sigma), 0.01 mM Alpha Amanitin (AAM, 23109-05-9, Caymen Chemical), or 10 µM RAD51 inhibitor BO2 (BO2, 1290541-46-6, Sigma) for 24 hours before imaging or fixation and immunofluorescence. VPA, AAM, and BO2 were dissolved in cell culture media.

### Transfection

Cells were transfected with the Neon transfection system (Cat.No.NEON1SK) and adapted from (Pabba et al., 2023) Cells were grown to 60-80% confluency. Cells were trypsonized as described above and then collected in a 15mL conical tube for centrifugation. Cells were centrifuged for 5 min at 3000g. Pellet was collected and resuspended with 150 ml of resuspension buffer R and transferred to a 1 ml microfuge tube. 1 ml of Cy3-dUTP (Enzo Biochem, # ENZ-42501) was then added and mixed with the transfection pipette. 100 ml of that solution was submerged into 2 ml of electrolytic buffer E2 and cells were electroporated (MEF [voltage-1005 V, width-35, pulses-2]). The electroporated cells were then plated into either 4 chamber or gridded single well dishes (Cellvis) with prewarmed, antibiotic free optiMEM (Fisher, Cat No. 31985062). After 3 hours media was changed to DMEM. Cells were incubated for 48 hours post transfection before imaging.

### Live cell time lapse fluorescence imaging

As previously described, we used either MEF NLS-GFP, HT1080 NLS-GFP, or HT1080 FUCCI stable cell lines to quantify nuclear shape and rupture (Berg *et al*., 2023; Pho *et al*., 2023). For tracking live cell chromatin motion CY3-dUTPs were transfected into cells. Images were acquired with Nikon Elements software on a Nikon Instruments Ti2-E microscope, Orca Fusion Gen III camera, Lumencor Aura III light engine, TMC CLeanBench air table, with 40x air objective (N.A 0.75, W.D. 0.66, MRH00401). Live cell time lapse imaging was possible using Nikon Perfect Focus System and Okolab heat, humidity, and CO2 stage top incubator (H301).

Cells were imaged in either 4 well cover glass dishes, 8 well cover glass chambers, or single well gridded dishes (Cellvis). For time lapse data, images were taken in 2-minute intervals during 3 hours with 6 fields of view for each condition single plane. For chromatin motion time lapses, mages were taken in 2.5-second intervals during 1 hour with 2 fields of view for each condition.

### Bleb count

MEF cells were treated with Hoechst 33342 at a dilution of 1:20,000 to 1:40,000 for 15 min before population imaging or imaged of NLS–GFP to analyze nuclear shape. Images were taken with nine fields of view for each condition, more than 100 nuclei were counted, and the percentage of cells showing blebbed nuclei calculated (>100) for each biological replicate (≥3). Nuclei were scored as blebbed if a protrusion 1 µm in diameter or larger was present, as previously outlined in Stephens et al. (2018).

### Nuclear rupture analysis

NLS–GFP MEFs were used for nuclear rupture analysis. Image stacks were analyzed using the NIS Elements AR Analysis software (Nikon). For each condition, total nuclei were counted at the first and last frame of the timelapse and averaged. Blebbed nuclei were counted as all nuclei that displayed a nuclear bleb at any time during the timelapse. Nuclear ruptures were determined by a >25% change in the NLS–GFP intensity in the cytoplasm to that in the nucleus using 5×5 µm boxes in each and with the background subtracted. Total nuclei showing a nuclear rupture were counted while differentiating between bleb-based ruptures and non-blebbed ruptures. Bleb-based ruptures were defined as nuclei showing a bleb prior to rupturing, whereas non-blebbed ruptures did not show a bleb on the nucleus. Rupture frequency was calculated by counting and averaging the number of ruptures for each rupturing nucleus.

Additionally, blebs formed during timelapse imaging were counted while differentiating between (1) blebs forming, rupturing and thereby stabilizing the bleb, and (2) blebs forming, but disappearing without showing a nuclear rupture, or blebs forming, rupturing but disappearing, and not being stabilized by the rupture. This analysis justifies counting the percentage of blebbed cells only at one or two timepoints as the dynamics of the formation of new blebs are captured. For each condition, three fields of views were analyzed for each replicate. Graphs were made to show the percentage of ruptures as the total number of ruptured nuclei by the total number of nuclei in each field of view.

### Nuclear height analysis

As previously described in (Pho *et al*., 2023), spinning disk confocal images were acquired with Nikon Elements software on a Nikon Instruments Ti2-E microscope with Crest V3 Spinning Disk Confocal, Orca Fusion Gen III camera, Lumencor Celesta light engine, TMC CleanBench air table, with Plan Apochromat Lambda 100× Oil Immersion Objective Lens (N.A. 1.45, W.D. 0.13 mm, F.O.V. 25 mm, MRD71970). Images were acquired at 0.2 µm z-slices with a total distance of 15 µm via 77 images. The center of one nucleus was selected at a time and a slice view was created. The fluorescence intensity profile for the z-slice was analyzed using a full-width half-maximum (FWHM) calculation giving the width of the fluorescence intensity peak corresponding to the height of the nucleus. Two measurements of nuclear height were averaged for each nucleus. Ten to 20 nuclei were measured for each condition.

### Micromanipulation force measurements

As previously described (Stephens *et al*., 2017; Currey *et al*., 2022), vimentin null (V−/−) MEFs were grown in a micromanipulation well to provide low-angle access via micropipettes. V−/− MEF nuclei were isolated from living cells via spray micropipette of the mild detergent Triton X-100 (0.05%) in PBS. The pull micropipette was used to grab the nucleus. The isolated nucleus was then grabbed at the opposite end with a precalibrated force micropipette and suspended in preparation for force-extension measurements. The pull pipette was moved at 50 nm/s to provide a 5-6 µm extension to the nucleus. Nucleus extension (µm) was measured as the change in distance between the pull and force micropipettes. Force (nN was measured as the deflection (µm) of the force micropipette multiplied by the premeasured bending modulus (1.2–2 nN/µm). The slope of the force versus extension plot provides the spring constant (nN/µm) for the short chromatin-dominated regime (<3 µm) and long-extension regime comprise of both chromatin and the lamin A-dominated strain-stiffening regime (>3 µm). Each nucleus was force vs. extension measured 3-4 times and averaged single nucleus spring constant measurement for each short and long regimes.

### Immunofluorescence

Cells were grown in eight-well dishes for 48 h prior to fixation. Treatment with drugs was done 24 h prior to fixation, which was done with a solution of 3.3% paraformaldehyde in 0.1% Triton X-100 in PBS for 15 min. Three washing steps were performed with PBS and the last one with PBS containing 0.06% Tween 20 (PBS-T). Cells were then blocked for 1 h at room temperature with a blocking reagent 2% BSA (Thermo Fisher Scientific) in PBS followed by staining with primary antibodies. The primary antibodies used were: anti-lamin A/C at 1:200 (Cell Signaling Technology, 4C11, mouse monoclonal antibody #4777), anti-H3K9ac at 1:500 (Cell Signaling Technology, C5B11, rabbit monoclonal antibody #9649), anti-pMLC2 at 1:200 (Cell Signaling Technology 3671, rabbit monoclonal antibody), CDT1 Antibody at 1:1,000 (Novus Biologicals, NBP1-58114-20ul, rabbit polyclonal antibody) anti-RNA pol II CTD repeat YSPTSPS (phospho S5/pSer5) at 1:1000 (Abcam, 4H8 – ChIP Grade, ab5408, mouse monoclonal antibody) and anti-RNA pol II CTD repeat YSPTSPS (phospho S2/pSer2) at 1:1000 (Abcam, ab5095, rabbit polyclonal antibody). Secondary antibodies were used at a 1:1000 dilution and include Alexa Fluor 488-, 555- or 647-conjugated goat anti-mouse or goat anti-Rabbit IgG (H+L), F(ab′)2 fragment (Cell Signaling Technology, 4408–4414). The nuclei were stained using Hoechst 33342 in PBS at a 1:40,000 dilution for at least 5 min. Cells were kept and imaged in PBS or mounted using Prolong Gold anti-fade mountant (Invitrogen, P36930) and incubated overnight in the dark. Fluorescence intensities were analyzed by measuring single nuclei as regions of interest and subtracting backgrounds as 30×30 pixel box with no cells. All single nucleus measurements were averaged over multiple fields of view to provide a single average intensity measurement for each experiment, for which there were at least three replicates. For comparison, all intensities were normalized with the mean value for untreated nuclei. Data was displayed as a Superplot using provided template (Lord *et al*., 2020).

### RNA labeling

RNA labeling was accomplished using Click-iT RNA Alexa Fluor 594 Imaging Kit (Invitrogen, C10330) as previously described (Jao and Salic, 2008). Cells were plated in eight-well plates (Cellvis, C8-1.5H-N), and grown and left untreated or treated with VPA and/or α-amanitin. EU was added at a final concentration of 1 mM to cells and incubated for 1 h. Cells were fixed and permeabilized as described above. After two PBS washes, the 500 µl formulation of the Click-iT reaction cocktail was added and allowed to incubate for 30 min in the dark. The reaction was terminated by removing the solution and washing the cells with the defined Click-iT reaction rinse buffer. The nuclei were stained using Hoechst 33342 in PBS at a 1:40,000 dilution for at least 5 min. The cells were rinsed two more times in PBS. Cells were kept and imaged in PBS or mounted using Prolong Gold anti-fade mountant and incubated overnight in the dark

### Chromatin Motion Tracking

Chromatin imaging was achieved through time-lapse imaging as previously described with time intervals of 2.5 sec for CY3-dUTP and 25 sec for NLS-GFP. Chromatin motion tracking was achieved using both Nikon NIS Elements the FIJI plugin Trackmate. Images were first cropped and aligned in Nikon, a 250 x 250 box was created around each nucleus. Trackmate settings were adapted from (Nozaki et al., 2017; Nozaki et al., 2023; Nagashima et al., 2019) and optimized for our imaging setup and transfection efficiency. Briefly, images were imported into FIJI and run through loG detect. This data was then exported to excel and run through an MSD macro (available upon request). MSD data was then plotted using excel. Where MSD is a set of *N* displacements *x*^2^ is given by 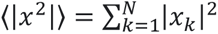.

## Abbreviations

NLS-GFP: nuclear localization signal green fluorescent protein
VPA: valproic acid
MEF: mouse embryonic fibroblast
EU: uridine analog 5-ethynyluridine
MSD: mean squared displacement
AAM: alpha amanitin
pMLC2: phosphorylated myosin light chain 2, UNT untreated

## Acknowledgements

We would like to thank Pierre Vidi, Edward J. Banigan, Tadasu Nozaki, and Kerry Bloom for their helpful and insightful discussions. We thank Erin Walsh for being a great lab mate and providing valuable feedback. We would like to thank John Orth for providing FUCCI cell lines. This work was supported by NIH NIGMS grant Maximizing Investigators’ Research Award R35GM154928, the Institute of Applied Life Sciences Midi-grant (190227), and by the Center for 3D Structure and Physics of the Genome 4DN2 grant (1UM1HG011536). The authors declare no competing interest.

## Data availability statement

The data are available in a public repository https://doi.org/10.6084/m9.figshare.29109695.v1

## Bibliography

Albiez, H, Cremer, M, Tiberi, C, Vecchio, L, Schermelleh, L, Dittrich, S, Küpper, K, Joffe, B, Thormeyer, T, von Hase, J, et al. (2006). Chromatin domains and the interchromatin compartment form structurally defined and functionally interacting nuclear networks. Chromosome Res 14, 707–733.

Banigan, EJ, Stephens, AD, and Marko, JF (2017). Mechanics and buckling of biopolymeric shells and cell nuclei. Biophys J 113, 1654–1663.

Barilla, D, Lee, BA, and Proudfoot, NJ (2001). Cleavage/polyadenylation factor IA associates with the carboxyl-terminal domain of RNA polymerase II in Saccharomyces cerevisiae. Proc Natl Acad Sci U S A 98, 445–450.

Belaghzal, H, Borrman, T, Stephens, AD, Lafontaine, DL, Venev, SV, Weng, Z, Marko, JF, and Dekker, J (2021). Liquid chromatin Hi-C characterizes compartment-dependent chromatin interaction dynamics. Nat Genet 53, 367–378.

Bensaude, O (2011). Inhibiting eukaryotic transcription: Which compound to choose? How to evaluate its activity? Transcription 2, 103–108.

Bercht Pfleghaar, K, Taimen, P, Butin-Israeli, V, Shimi, T, Langer-Freitag, S, Markaki, Y, Goldman, AE, Wehnert, M, and Goldman, RD (2015). Gene-rich chromosomal regions are preferentially localized in the lamin B deficient nuclear blebs of atypical progeria cells. Nucleus 6, 66–76.

Berg, HC (2018). Preface (1983). In: Random Walks in Biology, Princeton University Press, xi– 2.

Berg, IK, Currey, ML, Gupta, S, Berrada, Y, Nguyen, BV, Pho, M, Patteson, AE, Schwarz, JM, Banigan, EJ, and Stephens, AD (2023). Transcription inhibition suppresses nuclear blebbing and rupture independent of nuclear rigidity. J Cell Sci.

Bergamaschi, G, Biebricher, AS, Witt, H, Byfield, FJ, Seymonson, XMR, Storm, C, Janmey, PA, and Wuite, GJL (2024). Heterogeneous force response of chromatin in isolated nuclei. Cell Rep 43, 114852.

Bunner, S, Prince, K, Pujadas Liwag, EM, Eskndir, N, Srikrishna, K, McCarthy, AA, Kuklinski, A, Jackson, O, Pellegrino, P, Jagtap, S, et al. (2024). Decreased DNA density is a better indicator of a nuclear bleb than lamin B loss. J Cell Sci.

Bushweller, JH (2019). Targeting transcription factors in cancer - from undruggable to reality. Nat Rev Cancer 19, 611–624.

Chu, CG, Lang, N, Walsh, E, Zheng, MD, Manning, G, Shalin, K, Cunha, LM, Faucon, KE, Kam, N, Folan, SN, et al. (2025). Lamin B loss in nuclear blebs is rupture dependent while increased DNA damage is rupture independent.

Chu, F-Y, Clavijo, AS, Lee, S, and Zidovska, A (2024). Transcription-dependent mobility of single genes and genome-wide motions in live human cells. Nat Commun 15, 8879.

Currey, ML, Kandula, V, Biggs, R, Marko, JF, and Stephens, AD (2022). A versatile micromanipulation apparatus for biophysical assays of the cell nucleus. Cell Mol Bioeng.

Dhanasekaran, R, Deutzmann, A, Mahauad-Fernandez, WD, Hansen, AS, Gouw, AM, and Felsher, DW (2022). The MYC oncogene - the grand orchestrator of cancer growth and immune evasion. Nat Rev Clin Oncol 19, 23–36.

Ershov, D, Phan, M-S, Pylvänäinen, JW, Rigaud, SU, Le Blanc, L, Charles-Orszag, A, Conway, JRW, Laine, RF, Roy, NH, Bonazzi, D, et al. (2022). TrackMate 7: integrating state-of-the-art segmentation algorithms into tracking pipelines. Nat Methods 19, 829–832.

Galves, M, Sperber, M, Amer-Sarsour, F, Elkon, R, and Ashkenazi, A (2023). Transcriptional profiling of the response to starvation and fattening reveals differential regulation of autophagy genes in mammals. Proc Biol Sci 290, 20230407.

Göttlicher, M, Minucci, S, Zhu, P, Krämer, OH, Schimpf, A, Giavara, S, Sleeman, JP, Lo Coco, F, Nervi, C, Pelicci, PG, et al. (2001). Valproic acid defines a novel class of HDAC inhibitors inducing differentiation of transformed cells. EMBO J 20, 6969–6978.

Gu, B, Swigut, T, Spencley, A, Bauer, MR, Chung, M, Meyer, T, and Wysocka, J (2018). Transcription-coupled changes in nuclear mobility of mammalian cis-regulatory elements. Science 359, 1050–1055.

Halfmann, CT, Sears, RM, Katiyar, A, Busselman, BW, Aman, LK, Zhang, Q, O’Bryan, CS, Angelini, TE, Lele, TP, and Roux, KJ (2019). Repair of nuclear ruptures requires barrier-to-autointegration factor. J Cell Biol 218, 2136–2149.

Hatch, EM, and Hetzer, MW (2016). Nuclear envelope rupture is induced by actin-based nucleus confinement. J Cell Biol 215, 27–36.

Helfand, BT, Wang, Y, Pfleghaar, K, Shimi, T, Taimen, P, and Shumaker, DK (2012). Chromosomal regions associated with prostate cancer risk localize to lamin B-deficient microdomains and exhibit reduced gene transcription. J Pathol 226, 735–745.

Herbert, KM, Greenleaf, WJ, and Block, SM (2008). Single-molecule studies of RNA polymerase: motoring along. Annu Rev Biochem 77, 149–176.

Hobson, CM, Kern, M, O’Brien, ET, 3rd, Stephens, AD, Falvo, MR, and Superfine, R (2020). Correlating nuclear morphology and external force with combined atomic force microscopy and light sheet imaging separates roles of chromatin and lamin A/C in nuclear mechanics. Mol Biol Cell 31, 1788–1801.

Jao, CY, and Salic, A (2008). Exploring RNA transcription and turnover in vivo by using click chemistry. Proc Natl Acad Sci U S A 105, 15779–15784.

Kalinin, AA, Hou, X, Ade, AS, Fon, G-V, Meixner, W, Higgins, GA, Sexton, JZ, Wan, X, Dinov, ID, O’Meara, MJ, et al. (2021). Valproic acid-induced changes of 4D nuclear morphology in astrocyte cells. Mol Biol Cell 32, 1624–1633.

Kedinger, C, Gniazdowski, M, Mandel, JL, Jr, Gissinger, F, and Chambon, P (1970). α-Amanitin: A specific inhibitor of one of two DNA-dependent RNA polymerase activities from calf thymus. Biochem Biophys Res Commun 38, 165–171.

Keizer, VIP, Grosse-Holz, S, Woringer, M, Zambon, L, Aizel, K, Bongaerts, M, Delille, F, Kolar-Znika, L, Scolari, VF, Hoffmann, S, et al. (2022). Live-cell micromanipulation of a genomic locus reveals interphase chromatin mechanics. Science 377, 489–495.

Khatau, SB, Hale, CM, Stewart-Hutchinson, PJ, Patel, MS, Stewart, CL, Searson, PC, Hodzic, D, and Wirtz, D (2009). A perinuclear actin cap regulates nuclear shape. Proc Natl Acad Sci U S A 106, 19017–19022.

Kim, M, Krogan, NJ, Vasiljeva, L, Rando, OJ, Nedea, E, Greenblatt, JF, and Buratowski, S (2004). The yeast Rat1 exonuclease promotes transcription termination by RNA polymerase II. Nature 432, 517–522.

Kirkconnell, KS, Magnuson, B, Paulsen, MT, Lu, B, Bedi, K, and Ljungman, M (2017). Gene length as a biological timer to establish temporal transcriptional regulation. Cell Cycle 16, 259–270.

Kirkconnell, KS, Paulsen, MT, Magnuson, B, Bedi, K, and Ljungman, M (2016). Capturing the dynamic nascent transcriptome during acute cellular responses: The serum response. Biol Open 5, 837–847.

Komarnitsky, P, Cho, EJ, and Buratowski, S (2000). Different phosphorylated forms of RNA polymerase II and associated mRNA processing factors during transcription. Genes Dev 14, 2452–2460.

Lammerding, J, Fong, LG, Ji, JY, Reue, K, Stewart, CL, Young, SG, and Lee, RT (2006). Lamins A and C but not lamin B1 regulate nuclear mechanics. J Biol Chem 281, 25768–25780.

Liu, K, Patteson, AE, Banigan, EJ, and Schwarz, JM (2021). Dynamic nuclear structure emerges from chromatin cross-links and motors. Phys Rev Lett 126, 158101.

Locatelli, M, Lawrimore, J, Lin, H, Sanaullah, S, Seitz, C, Segall, D, Kefer, P, Salvador Moreno, N, Lietz, B, Anderson, R, et al. (2022). DNA damage reduces heterogeneity and coherence of chromatin motions. Proc Natl Acad Sci U S A 119, e2205166119.

Maarouf, A, Iqbal, F, Sanaullah, S, Locatelli, M, Atanasiu, AT, Kolbin, D, Hommais, C, Mühlemann, JK, Bonin, K, Bloom, K, et al. (2024). RAD51 regulates eukaryotic chromatin motions in the absence of DNA damage. Mol Biol Cell 35, ar136.

Manning, G, Li, A, Eskndir, N, Currey, M, and Stephens, AD (2025). Constitutive heterochromatin controls nuclear mechanics, morphology, and integrity through H3K9me3 mediated chromocenter compaction. Nucleus 16.

Mistriotis, P, Wisniewski, EO, Bera, K, Keys, J, Li, Y, Tuntithavornwat, S, Law, RA, Perez-Gonzalez, NA, Erdogmus, E, Zhang, Y, et al. (2019). Confinement hinders motility by inducing RhoA-mediated nuclear influx, volume expansion, and blebbing. J Cell Biol 218, 4093–4111.

Murrell, M, Oakes, PW, Lenz, M, and Gardel, ML (2015). Forcing cells into shape: the mechanics of actomyosin contractility. Nat Rev Mol Cell Biol 16, 486–498.

Nagashima, R, Hibino, K, Ashwin, SS, Babokhov, M, Fujishiro, S, Imai, R, Nozaki, T, Tamura, S, Tani, T, Kimura, H, et al. (2019). Single nucleosome imaging reveals loose genome chromatin networks via active RNA polymerase II. J Cell Biol 218, 1511–1530.

Nava, MM, Miroshnikova, YA, Biggs, LC, Whitefield, DB, Metge, F, Boucas, J, Vihinen, H, Jokitalo, E, Li, X, García Arcos, JM, et al. (2020). Heterochromatin-driven nuclear softening protects the genome against mechanical stress-induced damage. Cell 181, 800–817.e22.

Nozaki, T, Shinkai, S, Ide, S, Higashi, K, Tamura, S, Shimazoe, MA, Nakagawa, M, Suzuki, Y, Okada, Y, Sasai, M, et al. (2023). Condensed but liquid-like domain organization of active chromatin regions in living human cells. Sci Adv 9, eadf1488.

Pabba, MK, Ritter, C, Chagin, VO, Meyer, J, Celikay, K, Stear, JH, Loerke, D, Kolobynina, K, Prorok, P, Schmid, AK, et al. (2023). Replisome loading reduces chromatin motion independent of DNA synthesis. Elife 12.

Pho, M, Berrada, Y, Gunda, A, Lavallee, A, Chiu, K, Padam, A, Currey, ML, and Stephens, AD (2023). Actin contraction controls nuclear blebbing and rupture independent of actin confinement. Mol Biol Cell, mbcE23070292.

Pho, M, Berrada, Y, Gunda, A, and Stephens, AD (2024). Nuclear shape is affected differentially by loss of lamin A, lamin C, or both lamin A and C. MicroPubl Biol 2024.

Pujadas Liwag, EM, Acosta, N, Almassalha, LM, Su, YP, Gong, R, Kanemaki, MT, Stephens, AD, and Backman, V (2025). Nuclear blebs are associated with destabilized chromatin-packing domains. J Cell Sci 138.

Schwer, B, and Shuman, S (2011). Deciphering the RNA polymerase II CTD code in fission yeast. Mol Cell 43, 311–318.

Shaban, HA, Barth, R, and Bystricky, K (2018). Formation of correlated chromatin domains at nanoscale dynamic resolution during transcription. Nucleic Acids Res 46, e77–e77.

Shaban, HA, Barth, R, Recoules, L, and Bystricky, K (2020). Hi-D: nanoscale mapping of nuclear dynamics in single living cells. Genome Biol 21, 95.

Shimamoto, Y, Tamura, S, Masumoto, H, and Maeshima, K (2017). Nucleosome-nucleosome interactions via histone tails and linker DNA regulate nuclear rigidity. Mol Biol Cell 28, 1580–1589.

Stephens, AD, Banigan, EJ, Adam, SA, Goldman, RD, and Marko, JF (2017). Chromatin and lamin A determine two different mechanical response regimes of the cell nucleus. Mol Biol Cell 28, 1984–1996.

Stephens, AD, Banigan, EJ, and Marko, JF (2019a). Chromatin’s physical properties shape the nucleus and its functions. Curr Opin Cell Biol 58, 76–84.

Stephens, AD, Liu, PZ, Banigan, EJ, Almassalha, LM, Backman, V, Adam, SA, Goldman, RD, and Marko, JF (2018). Chromatin histone modifications and rigidity affect nuclear morphology independent of lamins. Mol Biol Cell 29, 220–233.

Stephens, AD, Liu, PZ, Kandula, V, Chen, H, Almassalha, LM, Herman, C, Backman, V, O’Halloran, T, Adam, SA, Goldman, RD, et al. (2019b). Physicochemical mechanotransduction alters nuclear shape and mechanics via heterochromatin formation. Mol Biol Cell 30, 2320–2330.

Strickfaden, H, Tolsma, TO, Sharma, A, Underhill, DA, Hansen, JC, and Hendzel, MJ (2020). Condensed chromatin behaves like a solid on the mesoscale in vitro and in living cells. Cell 183, 1772–1784.e13.

Strom, AR, Biggs, RJ, Banigan, EJ, Wang, X, Chiu, K, Herman, C, Collado, J, Yue, F, Ritland Politz, JC, Tait, LJ, et al. (2021). HP1α is a chromatin crosslinker that controls nuclear and mitotic chromosome mechanics. Elife 10.

Tan, MHE, Li, J, Xu, HE, Melcher, K, and Yong, E-L (2015). Androgen receptor: structure, role in prostate cancer and drug discovery. Acta Pharmacol Sin 36, 3–23.

Tinevez, J-Y, Perry, N, Schindelin, J, Hoopes, GM, Reynolds, GD, Laplantine, E, Bednarek, SY, Shorte, SL, and Eliceiri, KW (2017). TrackMate: An open and extensible platform for single-particle tracking. Methods 115, 80–90.

Tullai, JW, Schaffer, ME, Mullenbrock, S, Sholder, G, Kasif, S, and Cooper, GM (2007). Immediate-early and delayed primary response genes are distinct in function and genomic architecture. J Biol Chem 282, 23981–23995.

Vahabikashi, A, Sivagurunathan, S, Nicdao, FAS, Han, YL, Park, CY, Kittisopikul, M, Wong, X, Tran, JR, Gundersen, GG, Reddy, KL, et al. (2022). Nuclear lamin isoforms differentially contribute to LINC complex-dependent nucleocytoskeletal coupling and whole-cell mechanics. Proc Natl Acad Sci U S A 119, e2121816119.

Vargas, JD, Hatch, EM, Anderson, DJ, and Hetzer, MW (2012). Transient nuclear envelope rupturing during interphase in human cancer cells. Nucleus 3, 88–100.

Williams, JF, Surovtsev, IV, Schreiner, SM, Chen, Z, Raiymbek, G, Nguyen, H, Hu, Y, Biteen, JS, Mochrie, SGJ, Ragunathan, K, et al. (2024). The condensation of HP1α/Swi6 imparts nuclear stiffness. Cell Rep 43, 114373.

Zidovska, A, Weitz, DA, and Mitchison, TJ (2013). Micron-scale coherence in interphase chromatin dynamics. Proc Natl Acad Sci U S A 110, 15555–15560.

