## Supplemental Figures 1-4 for "Transcriptional activity generates chromatin motion that drives nuclear blebbing"

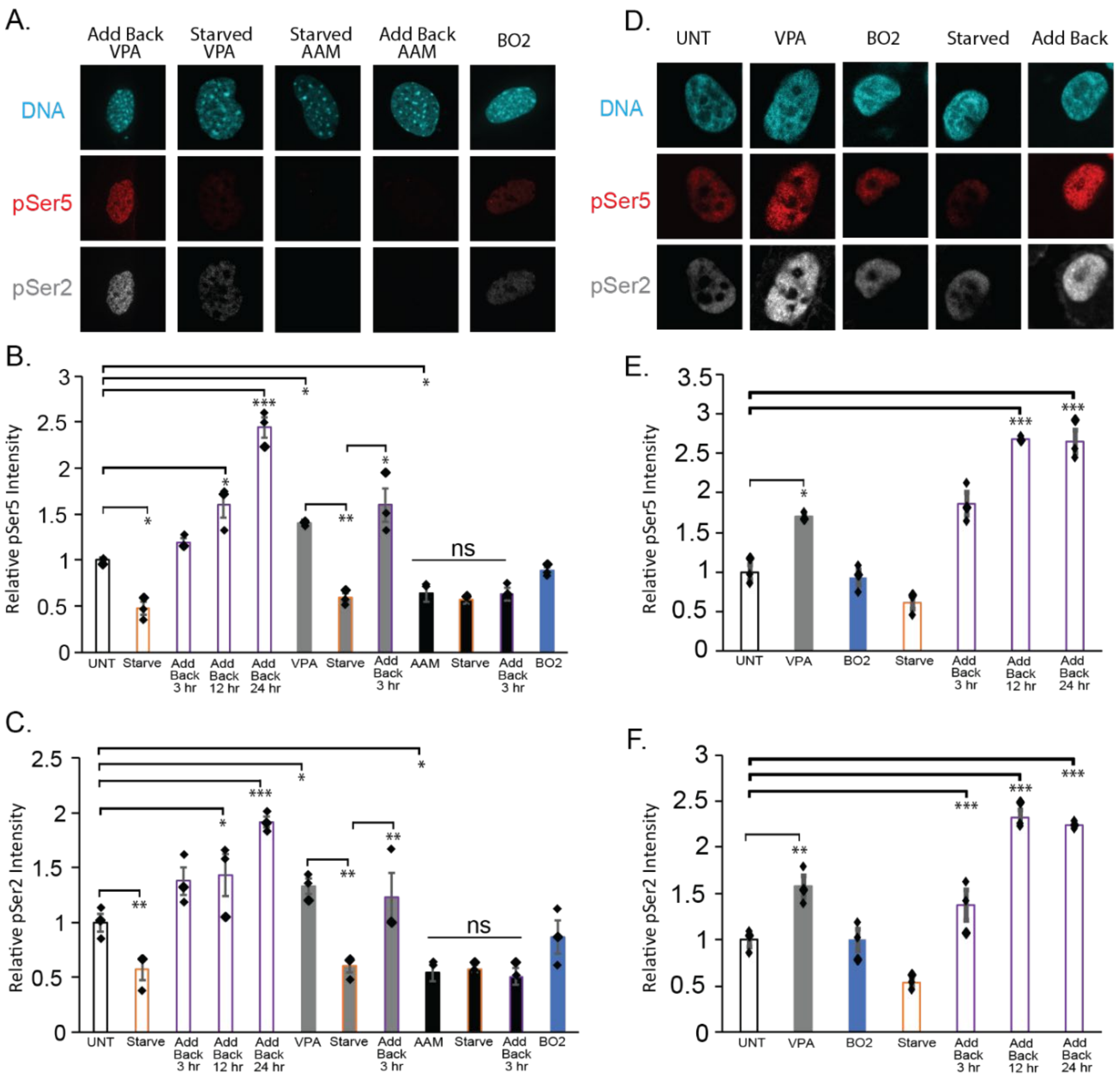


**Supplemental Figure 1. Quantification of active RNA Pol II markers pSer5 and pSer2.** (A) Example images of Hoechst staining for DNA, RNA Pol II markers pSer5 and pSer2. (B,C) Graphs of RNA Pol II pSer2 and p Ser5 in UNT, serum starved, serum add back (3, 12, and 14 hrs), VPA, Serum starved + VPA, serum add back + VPA, AAM, serum starved + AAM, serum add back + AAM, and BO2 in MEF WT cells. (D) Example images of Hoechst staining and RNA Pol II markers pSer2 and pSer5 in nuclear blebs of UNT, VPA, serum starved, and serum add back HT1080 WT cells. (E-F) Graphs of RNA Pol II fluorescent intensity pSer2 and pSer5 in HT1080 WT cells for UNT, VPA, BO2, serum starved, serum add back (3, 12, and 14 hrs). N=3 biological replicates with >20 nuclei for all graphs. Error bars represent standard error and statistical tests are one-way ANOVA with a post-hoc Tukey test, with significance denoted by *= p<0.05, **= p<0.01, and ***=p<0.001. Scale bar =10μm.

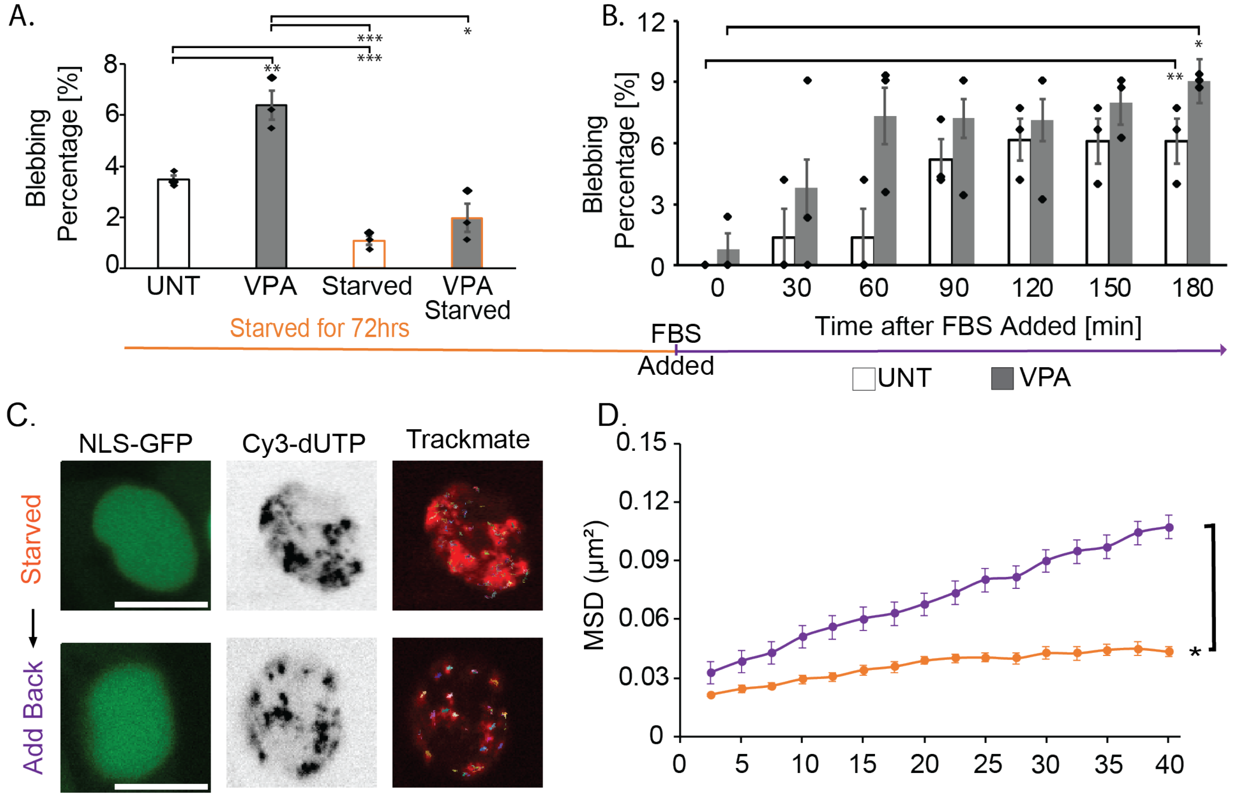


**Supplemental Figure 2. Nuclear blebbing percentages upon serum manipulation in HT1080 cells.** (A) Graph of blebbing percentage in HT1080 WT cells in conditions UNT, VPA, Serum starved, and VPA + serum starved. (B) Graph of change in blebbing percentage after 72-hour serum starvation, HT1080 cells are serum add back over 3 hours and blebbing is tracked in both UNT and VPA treated condition**.** N=3 biological replicates with >300 nuclei each. (C) Example images of wild type HT1080 nuclei with NLS-GFP and Cy3-dUTP used for tracking chromatin motion. Example Trackmate tracks for all measured chromatin foci domain motion for untreated serum starved (orange) and serum add back (purple). (D) Mean squared displacement (MSD) plots of HT1080 WT serum starved, and add back of >8 foci averaged per nucleus over 40 seconds, which is a quarter of full tracking of 150 seconds, n = 5 nuclei. Error bars represent standard error and statistical tests are one-way ANOVA with a post-hoc Tukey test (A-B) and two-tailed paired Student's t-tests (C-D), with significance denoted by *= $p<0.05$, **= $p<0.01$, and ***=$p<0.001$. Scale bar =10μm.

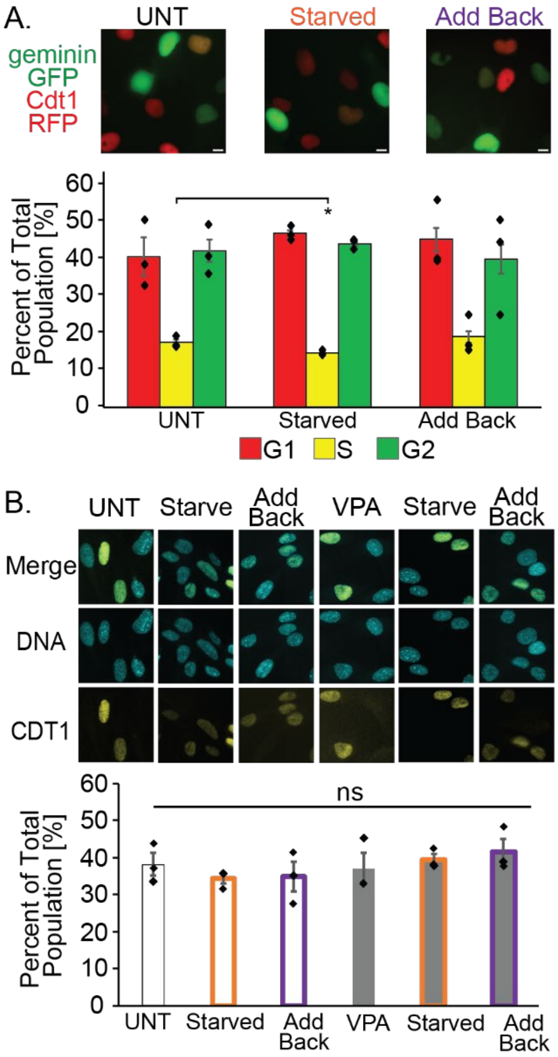


**Supplemental Figure 3. Cell cycle stage quantification upon serum manipulation. (**A) Example images of UNT, serum starved, and serum add back FUCCI HT1080 cells. Graph of % of cells in each of the interphase cell cycle stages (G1, S, and G2) for UNT, serum starved, and serum add back FUCCI HT1080 cells. (B)Example images of DNA stained with Hoechst and CDT1 for UNT, VPA, serum starved, serum add back, serum starved + VPA, and serum add back + VPA in MEF WT cells. Graph of CDT1 relative fluorescence CDT1 for UNT, VPA, serum starved, serum add back, serum starved + VPA, and serum add back + VPA in MEF WT cells. N=2 biological replicates with 6 technical replicates total with >20 nuclei per condition graphs. Error bars represent standard error and statistical tests are one-way ANOVA with a post-hoc Tukey test, with significance denoted by *= p<0.05, **= p<0.01, and ***=p<0.001. Scale bar =10 µm.

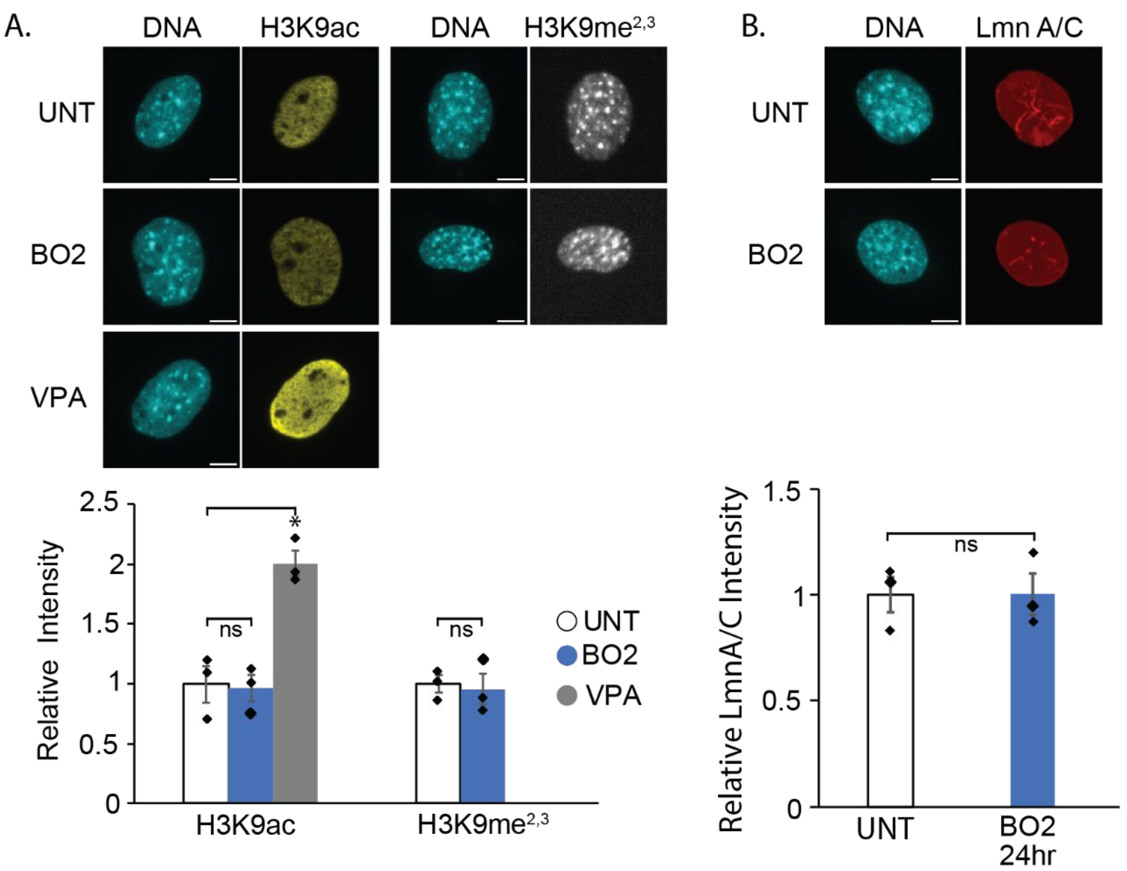


**Supplemental Figure 4. Eu-/heterochromatin and lamin levels measured in BO2-treated cells.** (A) Example images and graph of nuclei stained with Hoechst and H3K9ac euchromatin for UNT, VPA, and BO2, and H3K9me$^{2,3}$ heterochromatin marker for UNT and BO2 in MEF WT cells. (B) Example images and graph of nuclei stained for DAPI and lamin A/C for UNT and BO2-treated. Error bars represent standard error and statistical tests are two-tailed paired Student's t-test relative to UNT, with significance denoted by *= $p<0.05$, **= $p<0.01$, and ***=$p<0.001$. Scale bar =10μm.
